# Point-in-time evidence and cross-area clinical precedent anticipate clinical entry across 100 focal areas: retrospective validation of the Intangia triage layer

**DOI:** 10.64898/2026.08.19.745722

**Authors:** Thomas Elliott, Samuel Molnar, Gregory Peeters, Olivier Collart

**Author notes:** Correspondence Thomas Elliott, Intangia.

## Abstract

Early-opportunity teams face a combinatorial problem: once a focal target, mechanism or indication is fixed, the space of plausible partners runs to thousands of candidates per area. Intangia’s triage layer ranks that space from point-in-time evidence (how much literature, patent and clinical activity a candidate pairing has accumulated, and whether the partner already has clinical precedent in other contexts) so that review starts where clinical activity is most likely to begin next.

This preprint validates that capability retrospectively across 100 focal areas spanning drug targets, mechanisms and disease indications, replaying 24.12 million historically scored combination-years with every area scored by a model trained on the other 99 and never on itself. The headline is operational. At a twenty-partner review shortlist per focal area, the median area’s four-year first-alert precision is 0.234, against a matched random-ranker median of 0.008: roughly one in four shortlisted partners subsequently entered the focal clinical context within four years, about 38 times each area’s own background rate (95% CI 31 to 45). A panel-level permutation puts the result at empirical *p* ≤ 0.0005. Discrimination generalises: the full 13-feature specification reaches a median leave-one-focal-out ROC-AUC of 0.922 (95% CI 0.911 to 0.929), with no area below chance and all 100 areas beating their strongest count-based baseline. Shortlisted entrants are anticipated with a median observed lead of two years within the evaluation window, and three years (interquartile range one to five) once the window cap is removed and every realised entrant is counted.

The core ranking is carried by two interpretable signal families: cumulative co-occurrence counts and leave-one-area-out clinical precedent. Burst detection serves a complementary role: it supplies the time-stamped, source-specific momentum evidence attached to every recommendation (what is accelerating, and why now) rather than additional ranking power. A conditional view of the same landscape ranks candidates with no cross-area precedent against one another, enriched relative to matched random ranking, supporting a lower-yield emerging-opportunities capability. Worked replays in PD-1 combination immunotherapy and CTLA-4 make the output concrete: named partners entering the shortlist ahead of their first trials, each alert carrying the evidence dated at or before the historical cutoff under the study’s source timestamps.

**What this preprint validates:** Intangia has built a decision technology that detects where clinical activity is likely to emerge next. This preprint evaluates its first-stage triage layer: whether records timestamped at or before a historical scoring date can rank which focal-partner combinations will enter clinical trials within the following four years. Figure 1 shows where that layer sits in the wider decision pipeline, and what this study does and does not cover.

Figure 1.
Where the validated triage layer sits in the Intangia pipeline.
The upper band is what this preprint evaluates. The triage layer date-slices the evidence record of papers, patents and clinical trials using the source timestamps defined in Section 3.1, converts it into point-in-time signals, and ranks every eligible focal-partner pair with a single pooled cross-area model; cumulative co-occurrence counts and cross-area clinical precedent are the core ranking signals (Section 2.2). Burst detection runs as a parallel evidence lane rather than a driver of the core ranking: it identifies which source accelerated, when, and by how much, and that momentum evidence is attached to every recommendation (Section 2.4). The Invest and Watch tiers are a policy layer over the ranking that sets review depth and conviction bands, and no result in this paper depends on where their boundaries fall (Section 3.2). The lower band is how the shortlist is consumed in the product: agentic retrieval and assessment of current causal evidence for each shortlisted pair against a rubric the customer calibrates, and the decision-ready reports built from it. Those stages are shown for context only and none of them is evaluated here; the screenshots illustrate the interface rather than reporting results.

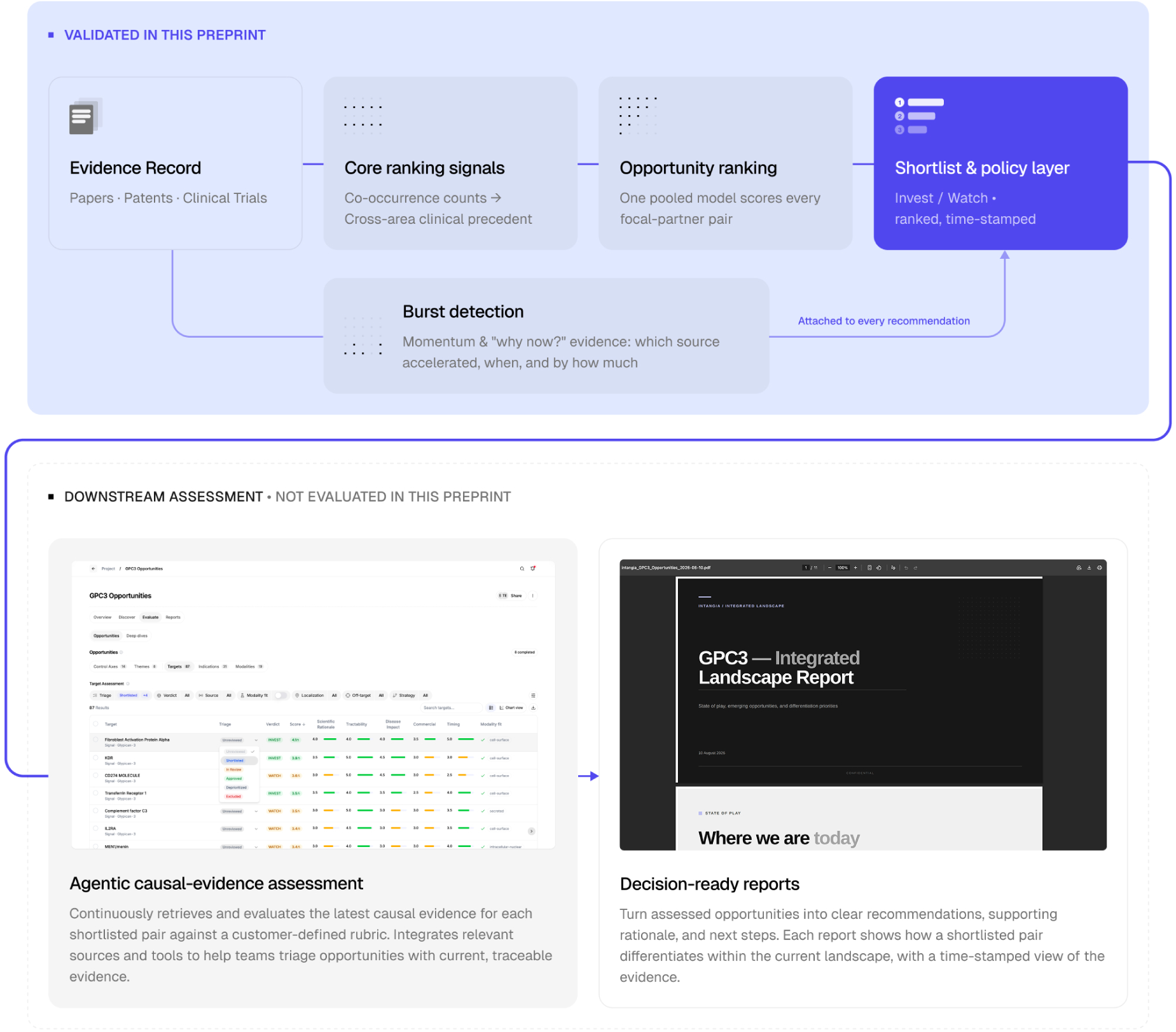

**Summary of validation results:** Every figure below is reported in full, with its protocol, in Section 2.

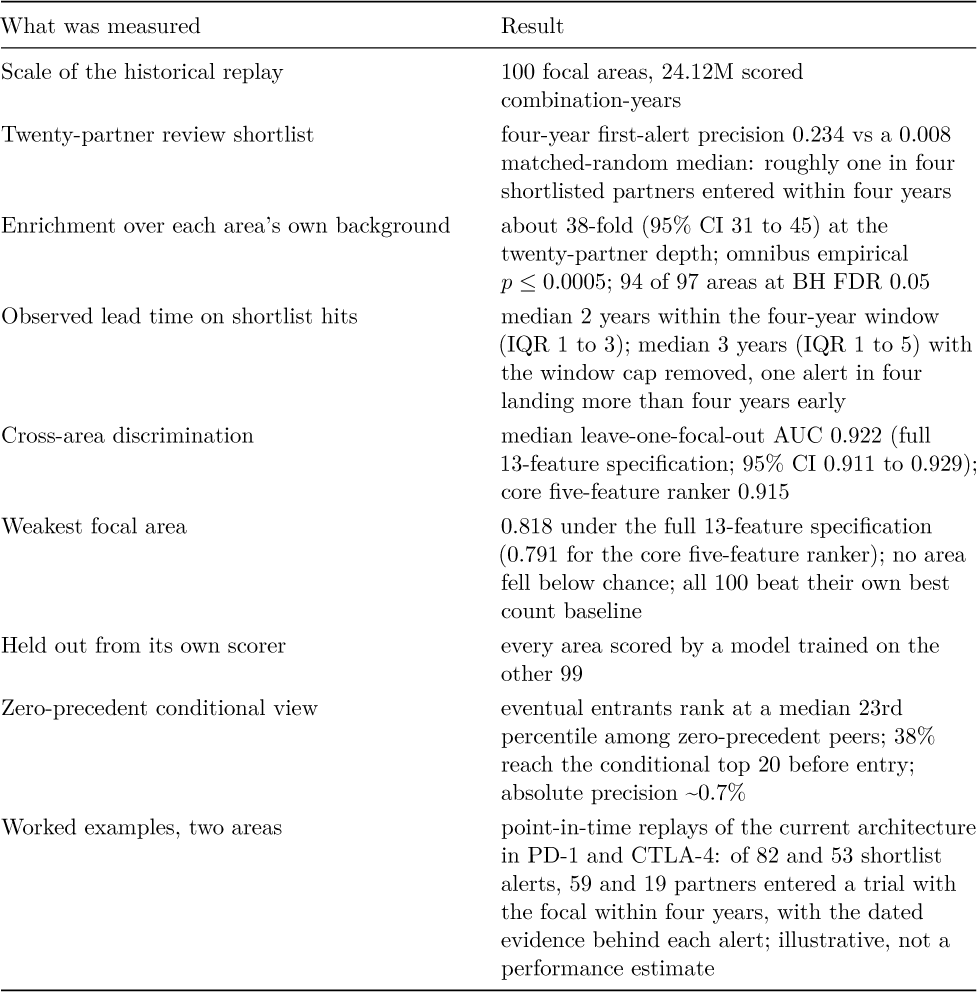

Every row above except the last is a validation result. The last is an illustration: PD-1 and CTLA-4 are worked examples showing what the system’s signals and lead times look like in familiar areas (Section 2.5). The evidence for how the layer performs in an area it has not seen comes from the 100-area held-out replay and the first-alert shortlist analysis.

The endpoint is first clinical entry: where development activity begins, not whether it succeeds. Biological causality, technical feasibility, clinical success and commercial value are assessed downstream and are outside the scope of this evaluation.

## 1. Introduction

The practical problem is combinatorial. Once a focal target is fixed, the number of plausible partner targets, pathways, indications, and mechanisms rapidly exceeds what can be evaluated experimentally or clinically. In immuno-oncology, this expansion has been especially visible around PD-1 and PD-L1 blockade, where many proposed combinations accumulated preclinical and commercial interest but only a subset became development programmes or entered trials [1–3]. The scale is substantial: by 2018 the immuno-oncology pipeline already spanned thousands of active combination trials and was expanding rapidly [42]. This combinatorial growth collides with a long-run decline in pharmaceutical research-and-development effiiciency (Eroom’s law, the observation that the inflation-adjusted cost of bringing a drug to market has risen steadily for decades [41]), which makes effiicient early triage of where to commit scarce experimental and analyst effort increasingly valuable. A triage method is therefore useful only if it can distinguish the small set of combinations likely to translate from a much larger set of incidental or non-productive co-occurrences.

Intangia addresses this problem as a staged decision-support process. The triage layer evaluated in this preprint narrows a broad evidence landscape into a ranked set of focal-partner opportunities for deeper scientific and strategic review. Its purpose is not to replace expert judgement, but to surface relevant programmes for earlier review, apply the same evidence criteria consistently across focal areas, and preserve the point-in-time rationale behind every ranking.

This problem sits within the lineage of literature-based discovery. Swanson’s work on undiscovered public knowledge showed that disconnected literatures can encode hypotheses not yet recognised explicitly by the field [4–6]. Subsequent work extended the ABC paradigm, introduced semantic and graph-based variants, and formalised time-sliced evaluation strategies for testing whether hypotheses generated from the past are borne out in future literature [7–11]. In parallel, biomedical knowledge graph methods have treated drug discovery as a link-prediction problem, integrating heterogeneous relations across drugs, targets, diseases, and mechanisms, including models for polypharmacy side effects using graph convolutional networks [12–16]. Text representation methods have also shown that embeddings trained on large corpora can capture latent scientific structure and, in some settings, anticipate later discoveries [17–20]. Separately, scientometric work on burst detection has provided tools for identifying periods of accelerating attention and emerging topics [21–24].

Two gaps remain. First, many discovery systems are evaluated with present-day data rather than under strict historical replay, which creates room for temporal leakage or for inadvertent use of evidence that was unavailable at the scoring date [25–27]. Second, rich machine-learning or graph models can produce accurate rankings while making it diffiicult to inspect what kind of record acceleration drove a prediction. The present study addresses these gaps with a triage layer whose inputs are deliberately simple and interpretable (point-in-time co-occurrence and burst signatures) even where the classifier that combines them is not, so that the discrimination claim rests on transparent features and a temporally controlled, point-in-time protocol rather than on model complexity.

### 1.1 Contributions

1. **Actionable shortlist performance** At a twenty-partner review depth, roughly one in four shortlisted partners subsequently entered the focal clinical context within four years: tens of times each area’s own background rate, and holding in almost every area on the panel. A landscape of thousands of candidates per area is reduced to a reviewable top twenty in which future entrants are concentrated, and they are typically flagged years before the trial (Section 2.1).
2. **Cross-area generalisation on an interpretable ranking signal** Under a temporally controlled historical replay of 100 focal areas, the model ranked future clinical entrants above the background landscape with strong and consistent discrimination: no area below chance, and every area beating its own best count-based baseline. Breadth spans oncology, immunology, metabolic disease, neuroscience, cardiovascular disease, haematology and rare disease. The signal is interpretable, with cumulative evidence counts and the partner’s leave-one-area-out clinical precedent carrying the ranking (Section 2.2).
3. **Time-resolved momentum evidence on every recommendation** Burst detection attaches to each flag the specific acceleration in the pre-cutoff record (which source moved, when, and by how much) so the rationale can be replayed at the date it was made and handed to expert review rather than accepted as a scalar score. It often reads as the earliest visible sign that something is beginning to move (Section 2.4).
4. **Worked demonstrations of the same architecture** Point-in-time replays in PD-1 combination immunotherapy and CTLA-4 show what the alerts looked like and what evidence stood behind each on the day it was made (Section 2.5). They are illustrative; the panel-wide analyses above carry the performance claims.

The first two contributions are the core validation; the latter two demonstrate how the technology is interpreted and consumed.

### Positioning

We did not identify a directly comparable historical-replay study combining the same focal-partner unit, first-clinical-entry endpoint and held-out-area design. The closest systems predict adjacent things (future citation of a paper by trials or guidelines, curated drug-disease links, target-disease advancement beyond phase 2, or topics that eventually yield approved drugs), and Table 8 (Section 4) places this work among them on unit, endpoint, temporal control and reported performance.

## 2. Results

**Box 1**. Units of analysis.

The evaluation is reported at several levels; the terms below are used throughout. Two of them are deliberately generic because the method does not distinguish the kind of entity involved: a *focal area* is the anchor concept a triage is run around, which may be a biological target, a mechanism of action, or a disease indication; a *partner* is a second concept found co-occurring with it, again a target, mechanism, or indication. A “combination” in this paper therefore means any such focal-partner pairing, not only a drug-drug combination therapy: it spans mechanism-by-indication pairings (the common monotherapy case) as well as target-by-target combinations (as in the PD-1 immunotherapy example). Where a statement is specific to one kind of entity, we say so.

| Term | Definition |
| --- | --- |
| Keyword row | One (focal area, co-occurring keyword, cutoff year) record; the native unit the model scores, and the level of the cross-area discrimination result. |
| Partner | The deduplicated unit of the operational analyses: keyword-level scores are collapsed to a single partner-level value by taking the maximum across resolved variants, and each partner can occupy only one shortlist position. |
| First alert | The first cutoff at which a partner enters the review shortlist (top $k$ unique partners of its cell). A partner counts once, at its first alert. |
| Invest / Watch flag | The point-in-time recommendation tiers presented to users: presentation bands over the ranking that set review depth and conviction. |

Worked example (PD-1 x JAK2). The pair (focal area = PD-1, co-occurring keyword = “Janus kinase 2”, cutoff year = 2015) is one *keyword row*, scored from the pre-2015 record. It is not the only row for this partner: “tyrosine-protein kinase JAK2” is a separate keyword row for the same target, and at the 2015 cutoff the two rank 40th and 1st in the area. Collapsing synonym variants by their best score makes them one *partner*, ranked 1st, so 2015 is that partner’s *first alert*. Its first registered PD-1 combination trial opened in 2016, within the four-year window of the alert, so under the first-alert protocol of Section 2.1 the alert counts as a hit with an observed lead of one year.

### 2.1 Shortlist performance: what a fixed review budget buys

At a twenty-partner review depth, the median area’s four-year first-alert precision is 0.234 against a matched random-ranker median of 0.008. Scores come from the core ranking signal, cumulative counts plus cross-area clinical precedent (the ablation in Section 2.2 is why that is the core model); the partner-level first-alert protocol, its matched null and the matching rules that leave 97 of the 100 areas evaluable are specified in Section 3.5.

That is roughly one in four shortlisted partners entering the focal clinical context within four years, in areas whose background rate makes the corresponding chance figure well under one in a hundred. The result is broad, with 94 of 97 areas individually surviving false-discovery control, and it degrades gracefully with depth, so a team can choose its own trade of precision against coverage along Table 1. Figure 2 shows the curve.

**Table 1.**
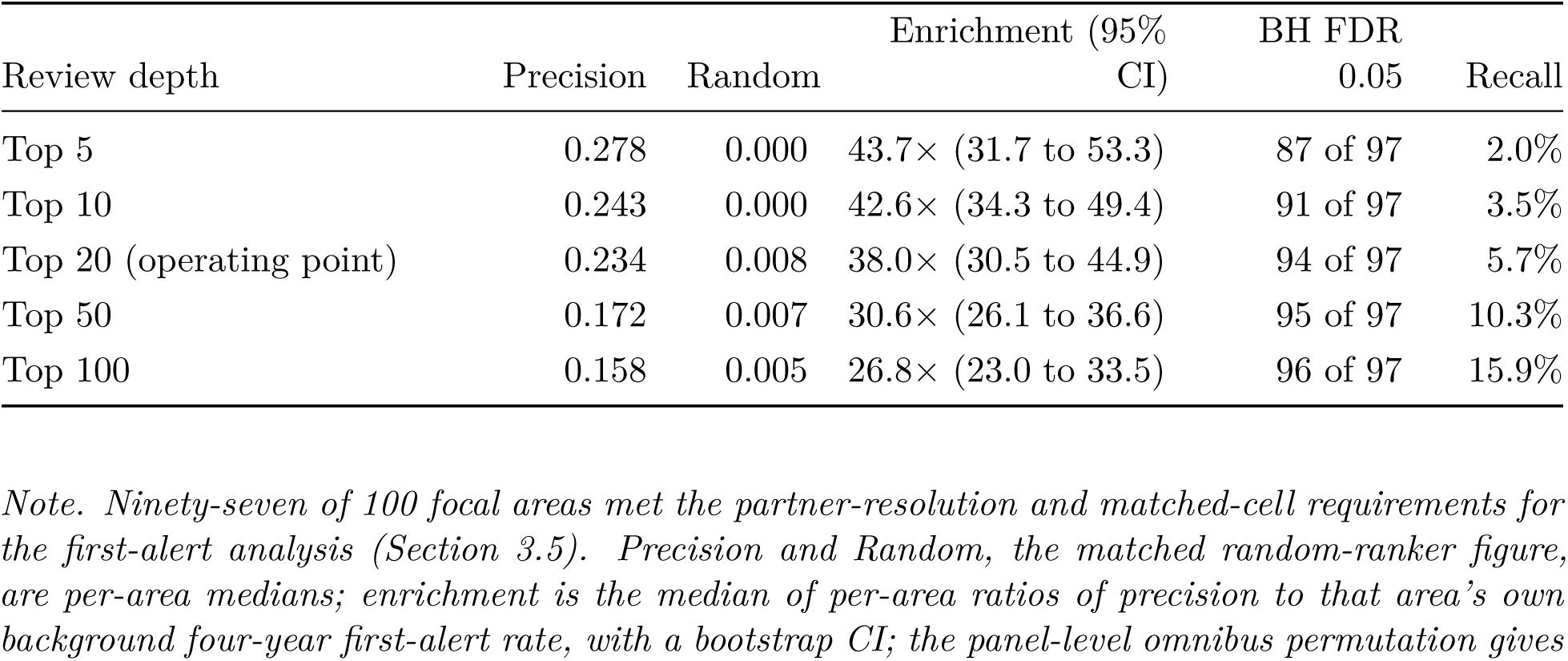
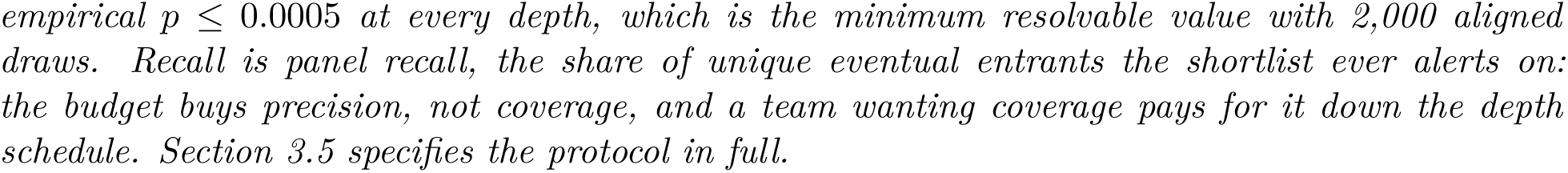
Four-year first-alert precision by review depth (medians across 97 areas).

| Review depth | Precision | Random | Enrichment (95% CI) | BH FDR 0.05 | Recall |
| --- | --- | --- | --- | --- | --- |
| Top 5 | 0.278 | 0.000 | 43.7× (31.7 to 53.3) | 87 of 97 | 2.0% |
| Top 10 | 0.243 | 0.000 | 42.6× (34.3 to 49.4) | 91 of 97 | 3.5% |
| Top 20 (operating point) | 0.234 | 0.008 | 38.0× (30.5 to 44.9) | 94 of 97 | 5.7% |
| Top 50 | 0.172 | 0.007 | 30.6× (26.1 to 36.6) | 95 of 97 | 10.3% |
| Top 100 | 0.158 | 0.005 | 26.8× (23.0 to 33.5) | 96 of 97 | 15.9% |

*Note. Ninety-seven of 100 focal areas met the partner-resolution and matched-cell requirements for the first-alert analysis (Section 3.5). Precision and Random, the matched random-ranker figure, are per-area medians; enrichment is the median of per-area ratios of precision to that area’s own background four-year first-alert rate, with a bootstrap CI; the panel-level omnibus permutation gives*
empirical $p \leq 0.0005$ at every depth, which is the minimum resolvable value with 2,000 aligned draws. Recall is panel recall, the share of unique eventual entrants the shortlist ever alerts on: the budget buys precision, not coverage, and a team wanting coverage pays for it down the depth schedule. Section 3.5 specifies the protocol in full.

**Figure 2.**
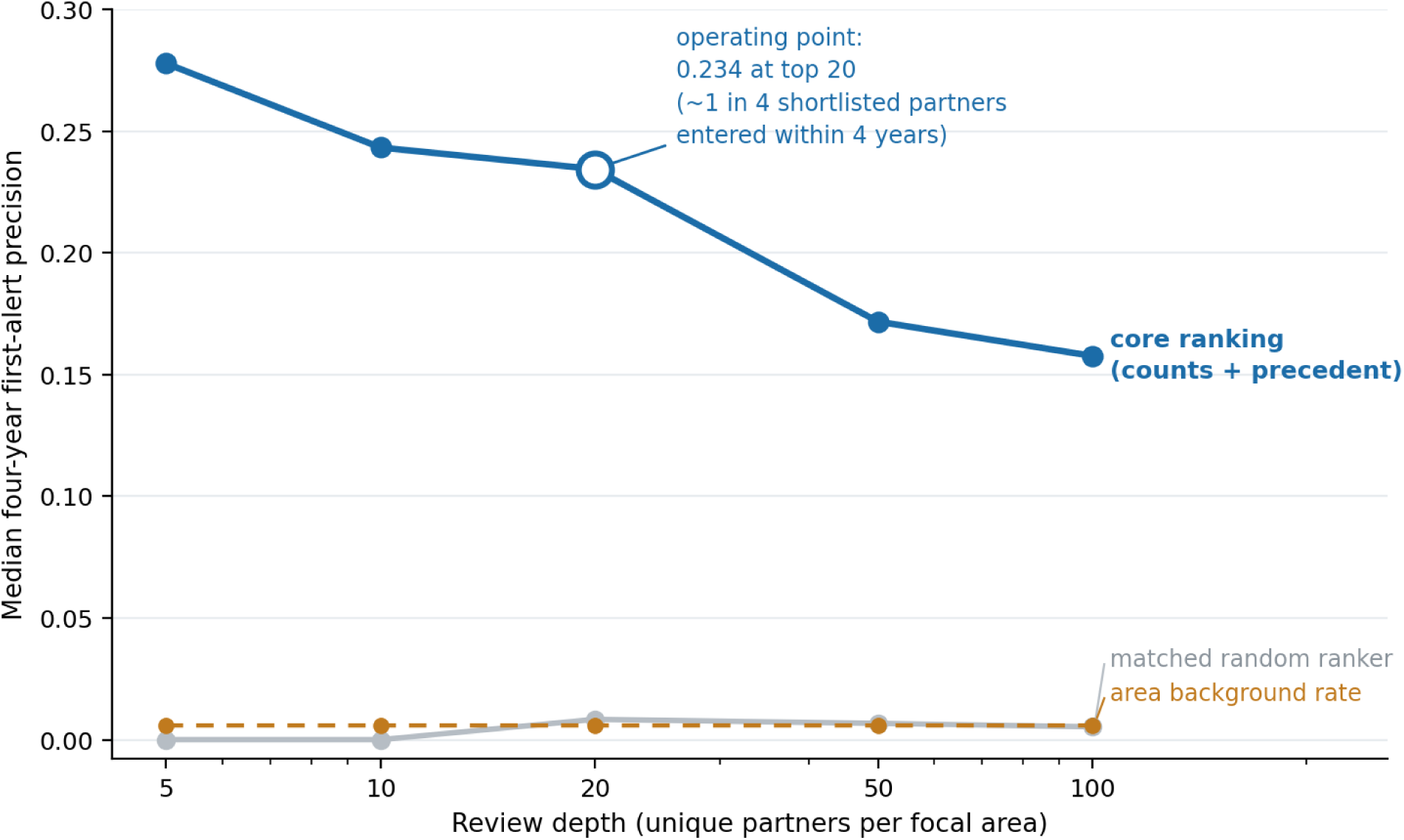
First-alert precision against review depth. Median per-area four-year first-alert precision of the core ranking (counts + cross-area clinical precedent) at shortlist depths of 5 to 100 unique partners per focal area, against the matched random-ranker median and the median per-area background rate under the identical protocol. The gap between the curve and its comparators is the operating value of the triage layer; the twenty-partner depth is the operating point used throughout this paper.

### How early the alerts land

Among true-positive first alerts at the twenty-partner depth, the observed interval from first alert to first trial has a median of 2 years (IQR 1 to 3), and the distribution is front-loaded: the probability that a shortlisted partner enters within one year of its first alert is 0.087, within two years 0.159, within three 0.215, and within four 0.262. These four figures are pooled over all alerts, whereas the 0.234 headline is the median across areas, so the four-year figure here is not the Table 1 value. The same median and IQR hold at every depth in Table 1. That figure is bounded by the endpoint itself: an alert counts as a hit only if entry follows within the four-year evaluation window, so longer head starts are scored as misses and can never appear in it. Removing the cap and counting every shortlisted partner that entered at any point in the observed record, the observed lead is a median of 3 years (IQR 1 to 5), and roughly one alert in four landed more than four years before the trial; a partner’s first entry into an area’s top 100, the earliest point the ranking takes notice, shows the same 3-year median (IQR 2 to 5). The uncapped figures are computed among realised entrants only, since partners alerted but not yet entered are censored, and long leads partly reflect persistent early promotion rather than timing skill (Section 6).

The twenty-partner depth was an existing practical review depth and is used as the primary operating point; the other depths characterise the precision-coverage curve. The headline does not hinge on the feature-set choice made in Section 2.2: the full thirteen-feature specification gives 0.203 at the same depth, roughly 25 times its matched null, under the identical protocol (Table 3).

### 2.2 Cross-area discrimination and the core ranking signal

Across 100 evaluable focal areas, the full 13-feature specification achieved, under the temporally controlled replay of Section 3.4, a median leave-one-focal-out ROC-AUC of 0.922 (95% confidence interval 0.911 to 0.929), a minimum focal-area AUC of 0.818 and a median focal-area precision-recall AUC of 0.080. Every focal-area AUC exceeded chance, and all 100 areas beat their own best count-based baseline. A relaxed-control sensitivity on the same feature set scores 0.933, a difference of 0.011 (Section 2.3), so the temporal controls cost a small, quantified amount and the stricter figure is the one reported throughout.

Leave-one-focal-out means the model scoring an area is trained on the other 99 and on cutoffs strictly earlier than the one being scored, so no area is fitted and evaluated on itself.

Area-level scores correlate negatively with how much clinical activity an area already carries (Pearson −0.65 against per-area positives; −0.50 on ranks): the broadest, most heavily co-published areas are the hardest, because co-occurrence there carries least information about any specific pairing, so a panel weighted toward specific targets scores higher than one weighted toward broad indications at identical model quality.

The breadth of this result is as important as the median. Strong discrimination was observed across oncology, immunology, metabolic disease, neuroscience, cardiovascular disease and additional haematology and rare-disease areas. The method therefore generalises across focal contexts rather than depending on one favourable biology.

**Table 2.**
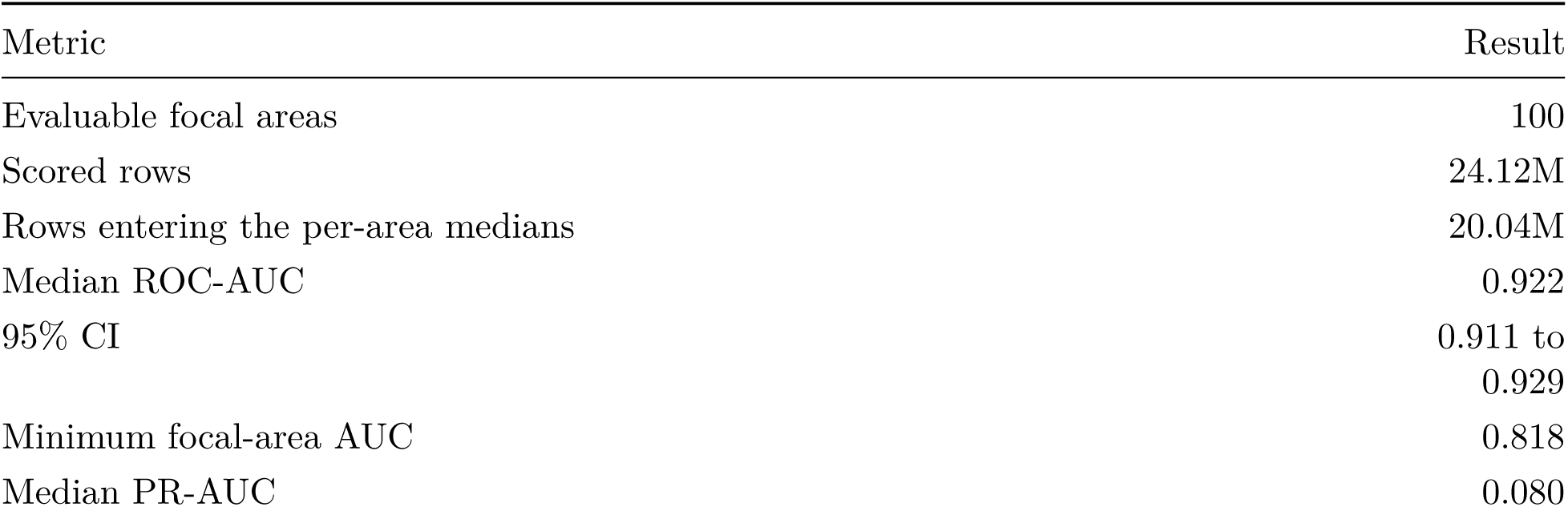
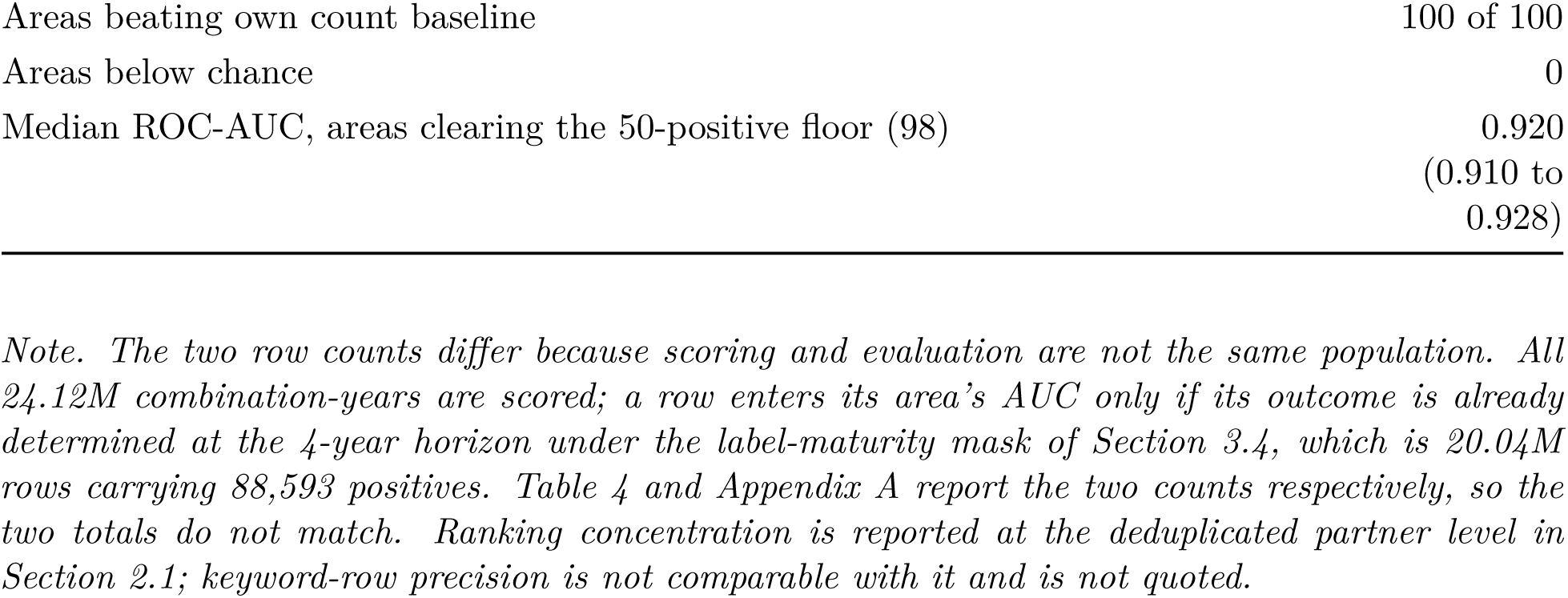
Primary discrimination performance (full 13-feature specification, temporally controlled replay).

| Metric | Result |
| --- | --- |
| Evaluable focal areas | 100 |
| Scored rows | 24.12M |
| Rows entering the per-area medians | 20.04M |
| Median ROC-AUC | 0.922 |
| 95% CI | 0.911 to 0.929 |
| Minimum focal-area AUC | 0.818 |
| Median PR-AUC | 0.080 |
| Areas beating own count baseline | 100 of 100 |
| Areas below chance | 0 |
| Median ROC-AUC, areas clearing the 50-positive floor (98) | 0.920<br>(0.910 to<br>0.928) |

*Note. The two row counts differ because scoring and evaluation are not the same population. All 24.12M combination-years are scored; a row enters its area’s AUC only if its outcome is already determined at the 4-year horizon under the label-maturity mask of Section 3.4, which is 20.04M rows carrying 88,593 positives. Table 4 and Appendix A report the two counts respectively, so the two totals do not match. Ranking concentration is reported at the deduplicated partner level in Section 2.1; keyword-row precision is not comparable with it and is not quoted.*

The summary statistics compress a full distribution, and the distribution itself is the more convincing object: Figure 3 plots the leave-one-focal-out AUC of every one of the 100 areas, ranked, with each area’s best count-based baseline marked for comparison. It makes the breadth claim directly verifiable rather than resting on the median and minimum alone.

**Figure 3.**
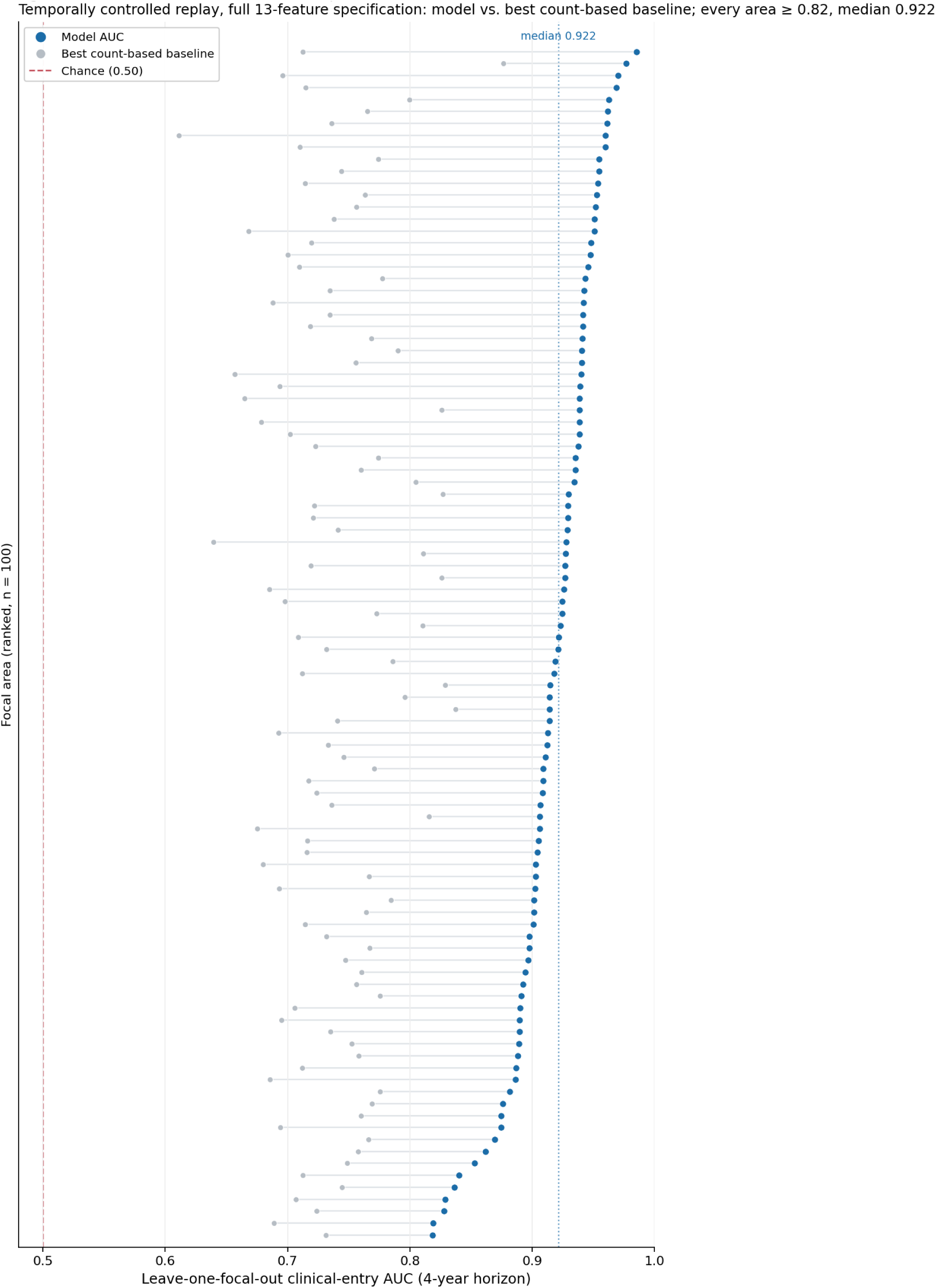
Per-area discrimination across the panel under the temporally controlled replay. Each point is one focal area’s leave-one-focal-out clinical-entry AUC (full 13-feature specification) at the 4-year horizon, ranked from lowest to highest; the paired open marker is that area’s best count-based bibliometric baseline. The dashed line at 0.50 is chance. Every area lands above 0.81 and none falls below chance, with a median of 0.922 and a minimum of 0.818. The spread is itself informative: the weakest areas are the broadest and most heavily studied concepts (Section 2.2), where co-occurrence carries least information about any specific pairing.

**Table 3.** Feature-group decomposition (temporally controlled replay, 100 areas, 4-year horizon).

| Feature set | Cols | Median AUC | Min AUC | Inverted | PR-AUC | Lift | P@20 |
| --- | --- | --- | --- | --- | --- | --- | --- |
| Counts + precedent (core ranker) | 5 | 0.915 | 0.791 | 0 | 0.069 | +0.173 | 0.234 |
| Full 13-feature specification | 13 | 0.922 | 0.818 | 0 | 0.080 | +0.180 | 0.203 |
| Precedent only | 2 | 0.883 | 0.757 | 0 | 0.068 | +0.148 | n/a |
| Full minus precedent | 11 | 0.749 | 0.406 | 1 | 0.016 | +0.007 | 0.036 |
| Counts only | 3 | 0.730 | 0.171 | 2 | 0.014 | +0.001 | n/a |
| Bursts only | 8 | 0.699 | 0.299 | 1 | 0.012 | −0.040 | n/a |

*Note.* Every row uses one panel, one protocol, one model class and identical held-out rows, asserted programmatically, so each line differs only in its feature set; the full specification is the core ranker’s five features plus the eight burst features. Min AUC is the worst per-focal value; “inverted” counts areas below 0.50; Lift is the median over each area’s best count-based baseline. P@20 is the twenty-partner first-alert precision of Section 2.1’s protocol, computed for the configurations evaluated operationally; the remaining cells were not run at the operational level.

The decomposition is clean, and it selects the core ranker. The full 13-feature specification achieved the highest global discrimination (median AUC 0.922), while the simpler five-feature counts-and-precedent configuration achieved the strongest operational shortlist performance and was selected as the core ranker. Cross-area partner precedent is the engine: two features alone reach a median AUC of 0.883, and the stronger of the two, the partner’s leave-one-area-out clinical-entry rate, reaches 0.887 entirely on its own. Adding the three cumulative count features brings the five-feature counts-and-precedent core ranker to 0.915, within 0.007 of the full thirteen-feature specification, and every configuration lacking precedent collapses toward the level of naive counting. At the operational endpoint the ordering tightens further: the core ranker matches or beats the full specification at every review depth (0.234 versus 0.203 at twenty partners), with identical recall and identical lead-time distributions. Counts + cross-area clinical precedent is therefore the validated core ranking signal. The burst features do not materially improve first-alert precision, recall or lead time once counts and precedent are present; their role (momentum, timing and the auditable evidence trail) is examined in Section 2.4.

### 2.3 Robustness

On the full 100-area panel the discrimination result holds up under a control-strictness sensitivity, a class-imbalance-aware metric, a label-permutation check and a feature ablation.

#### What the temporal controls cost

The temporally controlled replay imposes three restrictions beyond point-in-time feature construction: complete outcome-label maturation, a first-entry risk set, and leave-one-area-out precedent (Section 3.4). A relaxed-control sensitivity, which removes the maturation and at-risk restrictions while keeping point-in-time features, scores 0.933 on the identical feature set; imposing the full controls gives the 0.922 headline (Table 4). The controls therefore cost 0.011 median AUC, and the headline is reported under the stricter protocol throughout.

**Table 4.** Control-strictness sensitivity (100-area panel, 4-year horizon).

| Configuration | Rows | Base rate | Median AUC | Min AUC | PR-AUC |
| --- | --- | --- | --- | --- | --- |
| Relaxed-control sensitivity | 28.73M | 0.43% | 0.933 | 0.823 | 0.088 |
| Temporally controlled replay (headline) | 24.12M | 0.50% | 0.922 | 0.818 | 0.080 |

*Note.* Columns are median leave-one-focal-out AUC, per-focal minimum AUC, and median per-area precision-recall AUC. Base rate is pooled over all rows; the per-area median base rate, which the enrichment calculation below uses, is lower at 0.365% because the largest areas are also the densest. The relaxed-control row trains on every prior cutoff and scores all held-out rows; both rows use the full 13-feature specification, and the headline row additionally requires mature outcome windows on both sides of the split and restricts scoring to combinations still at risk of first entry. The 95% confidence interval on the headline is a cluster bootstrap resampling the 100 focal-level leave-one-focal-out units (the appropriate cluster for a median-across-focals statistic, and preferable to a row-level bootstrap given repeated combination-year observations): 0.911 to 0.929.

Because positives are rare and unevenly distributed, we report precision-recall AUC (average precision) alongside ROC-AUC. Computed per focal area as that area’s average precision divided by its own event prevalence and then medianed across areas (Section 3.5), enrichment is about 19-fold under the full 13-feature specification, with a 95% interval of 18 to 21. That figure is a median of per-area ratios, not the ratio of the median PR-AUC (0.080) to the median base rate (0.365%): the two medians are taken over different orderings of the areas, so dividing one by the other gives a different and larger number. This is the same rare-event concentration the shortlist analysis of Section 2.1 captures operationally, expressed here as a full precision-recall summary, and it confirms that the ROC-AUC headline does not rest on the extreme class balance.

A fixed-score permutation of the held-out labels, permuted within each focal area against the fitted predictions, produced a panel-median AUC of 0.500 over 300 repetitions (95% interval 0.498 to 0.502), confirming that the metric and the focal-area aggregation are centred at chance under label exchange.

The panel’s partner overlap is extensive: the median area shares 99.8% of its partners with at least one other area, with a minimum of 97.5%, so leaving out a focal area does not leave out its partners. That bounds the scope of the claim, which Section 6 states.

One further check bears on what the signal is made of. The feature ablation of Table 3 removes the partner-precedent features and drops the median from 0.922 to 0.749, isolating the burst-and-count contribution from the precedent contribution: the anticipatory signal rests on the partner’s prior clinical validation elsewhere, not on the focal pair’s own record. The event study of Section 2.4 makes the complementary temporal point: the clinical-source channel carries no information before a pair’s first trial exists, so the pre-trial signal lives entirely in the papers-and-patents record.

### 2.4 Burst analysis: momentum, timing and the evidence trail

Burst detection is the platform’s time-resolved channel: for every candidate pairing it identifies periods in which the paper, patent or clinical record accelerated, which source drove the acceleration, and when it began. Its validated role divides into three findings, kept deliberately separate.

#### The predictive finding

Adding the eight burst features to counts and precedent does not materially improve the ranking. The gain is +0.007 median AUC (Table 3), and at the operational endpoint there is no gain at all: first-alert precision at every review depth is equal or better without them, recall is identical, and the lead-time distributions are indistinguishable (Section 2.2). The same holds in the zero-precedent stratum, where the burst features’ contribution to discrimination is −0.001, so we do not claim that burst features materially improve prediction anywhere in the ranking. The scope of that statement should be read precisely. Candidate eligibility in this retrospective panel required at least one accepted historical burst. Within that burst-qualified candidate universe, burst-derived features did not materially improve ranking beyond cumulative counts and cross-area clinical precedent.

#### The product finding

What the burst channel supplies is the momentum evidence attached to every recommendation: a time-stamped, source-specific record of what is accelerating and why now, replayable at the date each flag was made. That evidence often arrives early. Among the zero-precedent eventual entrants surfaced into the conditional top twenty of Section 2.6 before entry, the first accepted burst preceded the first high-ranking alert in 84% (57 of 68). The count is taken on the trajectory reconstruction, which recovers pre-entry history for a wider set of entrants (223) than the scored sample behind the percentages of Section 2.6 (170), so its 68 surfaced entrants differ slightly from the roughly 65 implied there. The burst record tends to move before the candidate is promoted by the ranking, which is the early-warning, temporal-trigger role the channel plays in the product.

The same build-up is visible at population level. When the 72,174 translating combinations whose whole nine-year window falls inside the scored cutoff range are aligned at their first-trial year (T = 0), the mean text-only accepted-burst count rises steadily over the preceding years, 3.2-fold from T−6 to T−1 (0.0091 to 0.0295), in a gradual, monotonic climb rather than a spike, while the clinical-source channel only lights up after the first trial exists (Figure 4). Among combinations that eventually entered trials, the written record shows momentum gathering for years before the trial; the trial record itself carries no early information. The event study conditions on eventual entry, so it establishes temporal shape rather than discrimination.

**Figure 4.**
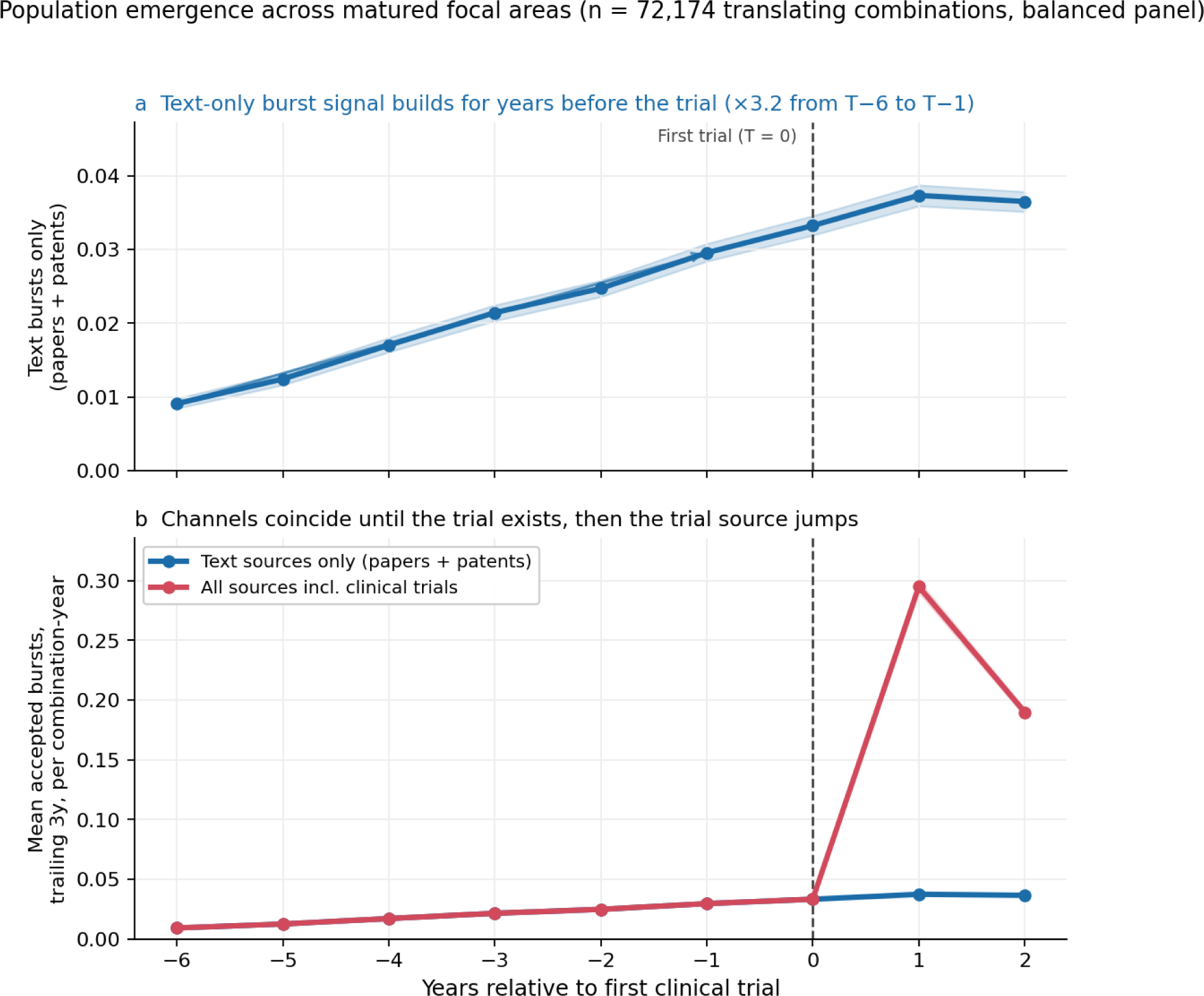
Literature and patent burst activity builds before first clinical entry. Each combination entering trials across the panel’s 100 focal areas is aligned at its first clinical-trial year (T = 0); curves are the mean number of accepted bursts in the trailing three years per combination-year at each relative year, with 95% confidence bands from a cluster bootstrap over whole combination trajectories. The panel is balanced: a combination is included only if every relative year it contributes falls inside its focal area’s scored cutoff range, so that a zero always means a scored cutoff at which no burst was accepted and never a year the panel does not reach. That admits 72,174 of the 155,192 translating combinations, and it keeps the comparison across relative years a statement about timing rather than about which combinations happen to be observable at each one. The two channels are the same statistic split by source scope and are therefore nested: a text-only channel (accepted bursts from papers and patents) and an all-source channel that additionally counts clinical-trial-source bursts. **(a)** The text-only channel on its own scale: burst activity rises 3.2-fold from T−6 to T−1, building gradually and monotonically before first trial. **(b)** Both channels on a common scale: they coincide until the first-trial year (a clinical-source burst cannot exist before the first trial) and diverge only at T+1 once trial-driven bursts appear. Event study over realised events: temporal shape, not predictive discrimination.

#### The exploratory finding

Within the zero-precedent pool, and after stratifying on fixed quartiles of underlying evidence volume, *recent* acceleration (a burst accepted within the last three years) and *multi-source* acceleration are associated with roughly two- to four-fold higher four-year entry rates in the mid- and high-volume strata (multi-source acceleration: 4.1 times the pool background), with the association absent among the most evidence-poor candidates. The direction is consistent but most strata are individually non-significant at these event counts, so this is a regime worth tracking as the record accumulates, not a claim of incremental predictive power and not the basis for a separate model.

One structural fact about the burst channel is worth recording for interpretation: candidate pairings enter the evidence panel through burst detection, so every scored candidate has at least one accepted burst in its history. “Has a burst” therefore carries no ranking information by construction; the informative dimensions are recency, repetition and source breadth, which are what the features encode and the exploratory analysis above stratifies.

### 2.5 Worked examples: point-in-time replays of the architecture in PD-1 and CTLA-4

PD-1 combination immunotherapy and CTLA-4 were selected post hoc as familiar illustrative examples from the panel. They are not used to estimate performance in an unseen focal area. This section replays the current triage architecture at each historical cutoff to show what its alerts looked like and what evidence stood behind them, and Sections 2.1 and 2.2 carry the performance claims.

In PD-1, the shortlist generated 82 first alerts across ten evaluable cutoffs, of which 59 (72%) entered the focal clinical context within four years, against an area background rate of 2.3%; the median observed lead is one year (IQR 1 to 2), short because in a crowded field entries follow the evidence quickly. In CTLA-4, 53 first alerts produced 19 entrants (36%) against a 1.2% background, with a median lead of three years (IQR 2 to 3). These are favourable areas on this metric: PD-1 ranks 3rd and CTLA-4 26th of the 97 evaluable areas on top-20 first-alert precision, and both background rates sit well above the panel median, which is exactly why the panel-wide figures of Section 2.1, not these two areas, are the performance estimate.

#### Coverage against depth

Twenty partners is a review budget, not a property of the ranking, which scores every burst-qualified candidate in the area (a median of 29,208 per cutoff in PD-1 and 7,843 in CTLA-4). At that budget the two areas cover a small share of what eventually happened: PD-1’s 82 alerts reach 4.8% of the 1,227 partners that eventually entered trials with it, and CTLA-4’s 53 reach 8.6% of its 222. Replaying the identical protocol deeper raises coverage to 16.0% and 38.7% at a hundred partners, 38.5% and 65.8% at five hundred, and 50.5% and 75.7% at a thousand. Precision decays as the list lengthens, from 72% to 28% in PD-1 and from 36% to 8% in CTLA-4, but a thousand-name list still runs about 12 and 7 times the area background rates quoted above, so the ordering carries information far below any depth a person would review by hand. Two boundaries keep this honest. Flagging every candidate in the area still reaches only 68.3% and 77.0% of eventual entrants, because the four-year conversion window binds regardless of depth; and precision at full depth is by construction the area’s own base rate, so a coverage figure means nothing unless the precision at that depth is quoted with it.

**Table 5.** Example alerts from the replayed shortlists. Selected hits with distinct partner identities (synonym variants collapsed), spanning target and indication partners; the complete alert lists, with every field below for all 135 first alerts, are available in the anonymised validation package described under Data and code availability.

| Area | Partner | First alert | First trial | Lead | Pair<br>evidence at<br>alert (papers<br>/ patents) | Partner<br>precedent<br>(areas) |
| --- | --- | --- | --- | --- | --- | --- |
| PD-1 | JAK3 | 2015 | 2018 | 3 y | 2 / 0 | 30 |
| PD-1 | Rheumatoid<br>arthritis | 2016 | 2019 | 3 y | 49 / 167 | 78 |
| PD-1 | TLR9 | 2014 | 2015 | 1 y | 16 / 33 | 51 |
| PD-1 | Adenosine<br>receptor<br>A2B | 2017 | 2018 | 1 y | 0 / 5 | 19 |
| CTLA-4 | CD47 | 2019 | 2023 | 4 y | 5 / 107 | 84 |
| CTLA-4 | AKT2 | 2014 | 2018 | 4 y | 3 / 2 | 64 |
| CTLA-4 | COX-2<br>(PTGS2) | 2014 | 2017 | 3 y | 3 / 9 | 82 |

*Note. “Pair evidence at alert” is the pair’s cumulative co-occurring paper and patent counts through the alert cutoff; “partner precedent” is the number of other focal areas in which the partner was already clinically active at that date: the two signal families the ranking runs on, shown at the moment the alert fired. CD47 is a recognisable case: alerted for CTLA-4 pairing in 2019 on a patent-heavy record (107 co-occurring patents against 5 papers) and 84 areas of prior clinical precedent, four years before the first registered trial pairing the two. A hit throughout is the first registered trial whose record pairs the two concepts, the panel’s uniform endpoint; where trial records name drugs rather than targets, that lexical match can lag or lead the true first co-administration. The candidate universe is deliberately unfiltered, with no self-reference exclusion and no cross-vocabulary synonym merge, so the full alert lists also contain same-axis entries (an anti-PD-L1 drug class alerted as a PD-1 partner) and duplicate name variants; this applies identically to every area in the panel-wide analysis of Section 2.1.*

Figure 5 draws every hit as its alert-to-entry interval. The picture a portfolio team should take from it is the product behaviour: a dated alert, carrying the evidence available to the replay under the study’s source-timestamp convention, lands a median of one to three years before the first registered trial, and the alert’s rationale (which sources were accumulating, and how much precedent the partner carried) is replayable per Table 5.

**Figure 5.**
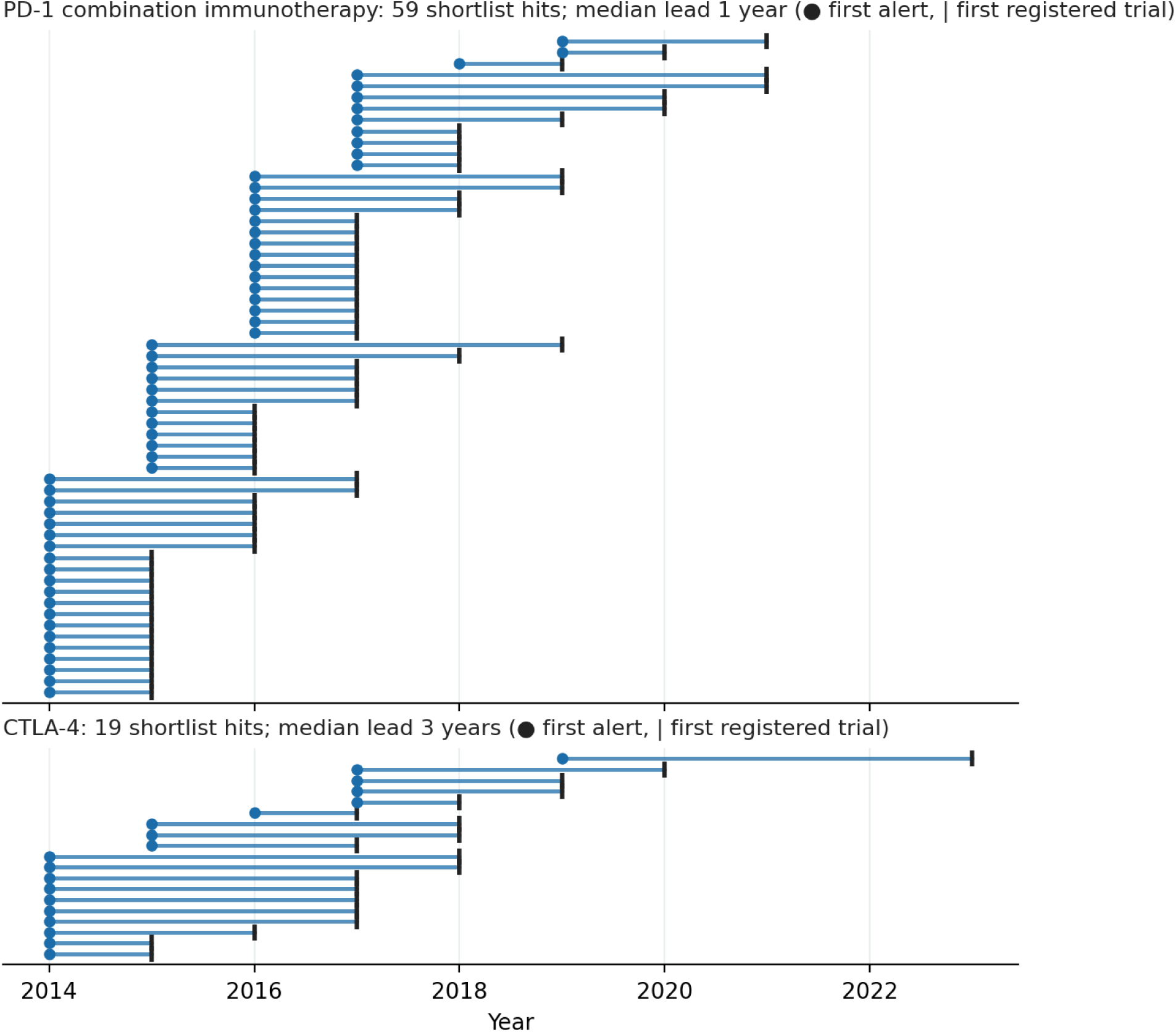
Alert-to-entry intervals in the two worked-example areas. One horizontal segment per true-positive first alert from the replayed twenty-partner shortlist: the dot is the first alert cutoff, the bar end is the partner’s first registered trial with the focal, and the segment length is the observed lead. PD-1: 59 hits, median lead one year; CTLA-4: 19 hits, median lead three years. Product demonstration under the protocol of Section 2.1; the panel-wide performance estimate is Table 1.

### 2.6 Emerging opportunities: ranking where no clinical precedent exists

Because cross-area clinical precedent carries much of the ranking, candidates whose partner has no clinical precedent in any other panel area sit at the bottom of the main list almost by construction. The product question is whether the same landscape supports a *conditional* view, ranking zero-precedent candidates against one another, and the answer is yes, with expectations set accordingly.

Among the zero-precedent pool (partners with a leave-one-area-out clinical-entry count of zero at the scoring cutoff), eventual entrants rank above the matched-random benchmark relative to their zero-precedent peers: the median eventual entrant sits at the 23rd percentile of its cell (chance is the 50th), and 38% of the 170 unique eventual-entrant area-partner pairs in the scored sample reach the conditional top twenty at least once before entry (25% reach the top ten; rank statistics computed on the full specification, and materially identical under the core model, whose conditional precision differs by under a tenth of a percentage point). Under the same first-alert protocol as Section 2.1, conditional top-twenty precision is about 0.7% against a matched within-pool random median of 0.3% and a pool background of 0.2%: roughly twice the matched null and three times background, but more than an order of magnitude below the precedented ranking of Table 1. For the entrants it surfaces, the conditional view is early: the median observed lead from first conditional alert to first trial is about three years. Within this pool the signal is carried by the evidence-volume features; the burst features do not add ranking power here either, though their momentum evidence often precedes the alert (Section 2.4).

The product reading is direct. The conditional view supports an emerging-opportunities lens: an alternative, lower-confidence ordering of the same landscape for teams scanning ahead of clinical precedent, presented with its own expectations rather than alongside the core shortlist’s hit rates. Two boundaries keep the claim honest: absolute precision is substantially lower than in the core ranking, and zero cross-area precedent within this panel is not the same thing as true first-in-class status, since a partner may hold precedent outside the panel’s scope, and first-in-class standing is a property of mechanism and indication, not of this panel’s precedent count.

## 3. Methods

### 3.1 Study design and data sources

This study evaluates the first-layer triage method of a broader target-prioritisation platform. The triage layer ingests three classes of record: scientific literature, patents, and clinical-trial records. The underlying co-occurrence tables are proprietary, but the study scope is explicit. Cross-area discrimination is evaluated on a panel of 100 focal areas spanning drug targets and disease indications, all 100 of which have an evaluable clinical leg (Table 6).

A fifty-positive reporting floor accompanies the panel: an area with very few positive outcomes yields a noisy AUC, so areas below fifty evaluated positives are identified rather than silently pooled. Two of the hundred fall below it, CACNA1H (the Cav3.2 calcium channel subunit) with 11 positives and RBX1 with 39, and both score above the median, so they add upward rather than downward noise. Applying the floor leaves 98 areas and moves the median AUC from 0.922 to 0.920 (95% CI 0.910 to 0.928), leaves the minimum unchanged at 0.818, and leaves every retained area beating its own count baseline. Because the effect is smaller than the width of the confidence interval, the tables in Section 2 are reported on all 100 areas, with the floor-applied median carried alongside in Table 2 and the two thin areas identified by their positive counts in Appendix A.

**Table 6.**
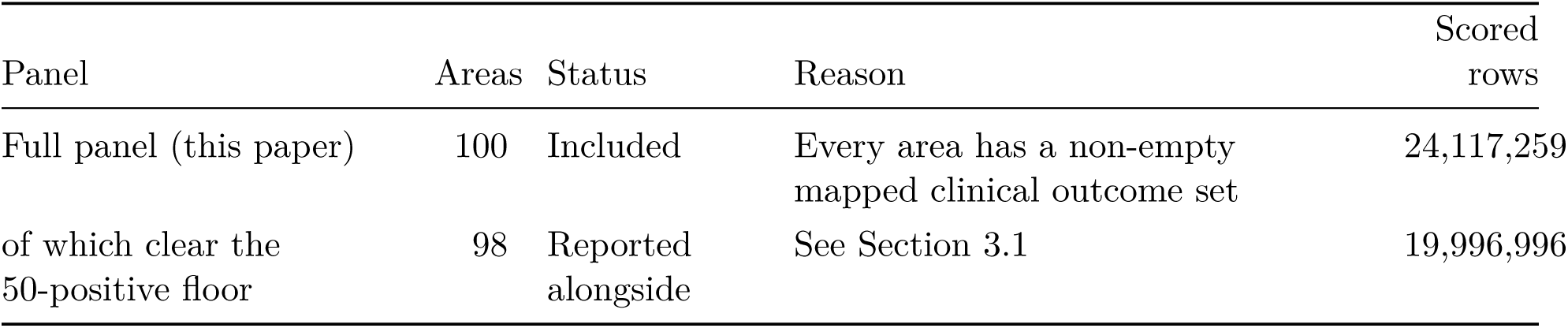
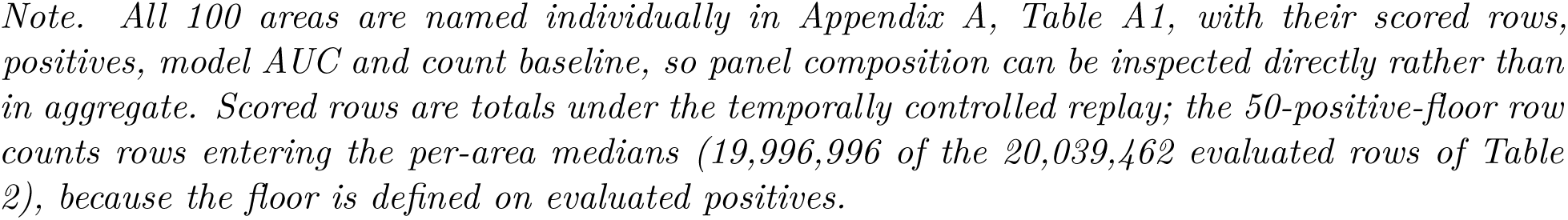
Panel composition.

| Panel | Areas | Status | Reason | Scored rows |
| --- | --- | --- | --- | --- |
| Full panel (this paper) | 100 | Included | Every area has a non-empty mapped clinical outcome set | 24,117,259 |
| of which clear the 50-positive floor | 98 | Reported alongside | See Section 3.1 | 19,996,996 |

*Note. All 100 areas are named individually in Appendix A, Table A1, with their scored rows, positives, model AUC and count baseline, so panel composition can be inspected directly rather than in aggregate. Scored rows are totals under the temporally controlled replay; the 50-positive-floor row counts rows entering the per-area medians (19,996,996 of the 20,039,462 evaluated rows of Table 2), because the floor is defined on evaluated positives.*

#### Temporal provenance

Every point-in-time claim in this paper rests on which date field assigns a record to a year, so we state them explicitly. The scientific literature is assigned to a year by publication date. Patents are assigned by patent publication date, not by priority or filing date, and are not deduplicated at patent-family level, so an invention published in several jurisdictions contributes one count per publication, spread across the years its family members published. Concept matching against these two corpora is performed upstream of this study by the indexing provider; what this study controls, and what every temporal claim below rests on, is that each record is dated by its publication date and that no record is re-indexed or re-dated here. The aggregate clinical series, which defines first clinical entry for the outcome label, the point-in-time in-trial indicator, and the clinical-source burst component, assigns a trial to its registration (first-submission) date, not its study-start date; the separate phase-resolved series maintained elsewhere in the platform are keyed to study start and are not used for the primary endpoint. Using the first-submission timestamp rather than study start generally produces an earlier entry date and therefore a shorter, more conservative estimated lead time, though retrospective registration can invert that ordering for individual trials. One consequence should be stated plainly. The clinical series is date-sliced using the first-submission timestamp recorded in the current registry extract. This reconstructs clinical-event timing, but it does not reproduce archived public registry availability at every historical date, and a recorded first-submission date does not by itself establish that the record was publicly observable on that date. The distinction matters because the clinical series drives the outcome label, the point-in-time in-trial indicator, the clinical-source burst component and the cross-area partner precedent that carries much of the discrimination. Papers and patents are not affected: both are sliced on publication date, which is availability. A first-posted-date sensitivity would close the gap and is listed in Section 7. Co-occurrence is measured at the document level: two concepts co-occur in a record when both are matched anywhere in that record’s indexed text, with no sentence- or passage-level proximity constraint, and each source-year figure is a count of distinct documents containing both rather than a count of mentions. Concept synonyms are folded onto the canonical concept before counting.

The analysis runs on a single extract of the co-occurrence tables generated on 14 July 2026. That extract carries no explicit corpus freeze date: each source is drawn up to its generation date, so coverage thins toward the final years of the window. This is precisely why label maturation is enforced per focal area against that area’s own observed clinical extent rather than against a single global freeze date (Section 3.4).

### 3.2 Feature engineering

Each candidate combination is represented by point-in-time features computed only from records dated at or before a chosen cutoff year Y, under the source timestamps set out in Section 3.1. The model uses an explicit, auditable feature vector in three groups.

*Burst and acceleration features* (eight) summarise the timing, intensity and breadth of joint activity: the number of bursts; the consensus score defined below; the count of accepted bursts in the trailing three years; a burst-acceleration ratio, recent accepted bursts divided by earlier accepted bursts; the number of text sources, papers and patents, contributing an accepted burst, valued 0 to 2; the number of distinct sources contributing any accepted burst including the trial record, valued 0 to 3; and two recency measures, the years elapsed since the combination’s first and since its most recent accepted burst.

*Count features* (three) and *partner-precedent features* (two) are defined below. The full evaluated feature vector therefore contains 13 features: eight burst and acceleration measures, three co-occurrence count measures, and two leave-one-area-out partner-precedent measures. The production core ranking uses the five count and precedent features (Section 2.2); the burst features are evaluated alongside them and carried in the product as the momentum-evidence channel (Section 2.4).

Because the primary endpoint is first clinical entry, combinations already in a trial at cutoff Y are excluded from the temporally controlled risk set, and the in-trial indicator, which records whether a combination is already in a trial as of Y, is not used as a predictor. It appears only inside the relaxed-control sensitivity, where it is point-in-time rather than a leak, because a combination already in a trial cannot be a *new* entry in the forward window.

*Count features* are the cumulative co-occurring paper and patent counts through Y and recent three-year paper activity. These same counts serve as the count-based baselines (Section 3.5).

*Precedent features* summarise the partner’s broader translation history as of Y. The first is partner recurrence, the number of distinct focal areas in which the partner is active by Y. The second is the partner prior *π*, its observed clinical-entry rate as of Y; that is, the fraction of focal areas in which the partner has already entered trials, shrunk toward the global base rate by a fixed pseudocount of 50. Writing *g* for the global base rate and counting distinct focal areas other than the one being scored, *π* = (entered+ 50*g*)/(present+ 50). Both counts exclude the scored area’s own contribution, so the feature is strictly leave-one-area-out, and *π* is built from point-in-time in-trial status rather than from any forward label.

Every feature above derives from unweighted co-occurrence counts and their dynamics. None is weighted by citation count, journal, or impact factor, so the signal reflects the volume and timing of joint activity rather than the prominence of the venues in which it appears. Because the burst features are evaluated as lift over the cumulative count baselines (Section 2.2), any effect attributable purely to high-visibility work being counted more would already be captured by the counts; the burst features are therefore evaluated net of that effect, and their measured incremental contribution is reported in Sections 2.2 and 2.4.

Burst detection runs exactly three complementary per-source detectors, independently on each source’s annual co-occurrence series: PELT changepoint detection in mean and variance, a derivative-based acceleration test, and a rolling z-score threshold (Table 7). Each detector is subject to a minimum-support constraint (a burst must rest on at least a minimum co-occurrence count to be accepted). Because three detectors are active, the consensus vote bonus, min(*a*/3, 1) × 0.5 for *a* detectors agreeing, saturates only at unanimous agreement. The hyperparameters are listed in Table 7; they were fixed a priori rather than tuned against the clinical-entry outcome, and no hyperparameter or detector threshold was optimised on the labels or on the held-out areas. The choice between the five-feature and thirteen-feature specifications was made on held-out operational precision, and both are reported throughout so the effect of that choice is visible (Table 3).

The annual series is smoothed with a three-year centred moving average before detection. The series is rebuilt at each cutoff and truncated at that cutoff, so the window at year Y spans only [Y−1, Y] and never reaches Y+1: the smoother is non-causal for interior years but stays entirely inside the information set dated at or before the scoring cutoff.

Two recency adjustments apply to cutoff years 2024 and later, where a genuine burst has had less time to accumulate the support the standard thresholds assume: the derivative detector’s z-threshold is multiplied by 0.9, and the rolling detector inflates the computed z-score by 1.1 while trimming its baseline to the last three points. Both relax the acceptance test. Neither is exercised anywhere in this study, because all cutoffs fall between 2010 and 2023; they are documented as deployment behaviour only and are omitted from Table 7.

**Table 7.** Burst-detection and consensus hyperparameters.

| Component | Parameter | Value |
| --- | --- | --- |
| Series construction | smoothing window (centred moving average) | 3 years |
|  | minimum non-zero years for inclusion | 2 |
| PELT changepoint (mean + variance) | penalty | MBIC, $4 \cdot \log(n)$ |
|  | minimum segment length | 3 years |
|  | acceptance | positive mean shift only;<br>co-occurrence count of at least 1 |
| Derivative / acceleration | z-threshold on smoothed first difference | 1.5 |
| Rolling z-score | trailing baseline window | 3 years |
|  | z-threshold | 2.0 |
| Consensus | detector weights (PELT / rolling / derivative) | 1.2 / 0.8 / 1.0 |
| | consensus score | weighted mean of detector scores $\times (1 + \text{vote bonus})$ |
| | vote bonus | $\min(\text{detectors agreeing} / 3, 1) \times 0.5$ |
|  | source acceptance (consensus score / effect size / detectors agreeing): clinical | 2.0 / 0.005 / 1 |
|  | patents | 5.0 / 0.01 / 2 |
|  | papers | 25.0 / 0.05 / 2 |

For presentation, the platform assigns each candidate a discrete tier at every cutoff as a policy layer over the ranking: Invest is the top 5% of the focal-area-cutoff cell by point-in-time score and Watch the next 20%. The tiers set review depth and conviction bands for users; no result in this paper depends on the tier boundaries, and the operational evaluation uses explicit review depths instead (Section 3.5).

### 3.3 Model choice

One pooled model family carries the study: a gradient-boosted tree ensemble (histogram gradient boosting) trained across areas, which with enough areas to train on generalises to areas it has not seen. The core ranker and the full 13-feature specification are this same model class fitted under the identical protocol; they differ only in which feature columns they receive (Table 3). This family is behind the primary result and all robustness checks (Sections 2.1 and 2.2). The discrimination claim rests on interpretable, point-in-time input features and a temporally controlled protocol rather than on the classifier: Section 2.2 decomposes the contribution of each feature group, so what the model exploits is transparent even though the classifier itself is not linear.

One recommender carries every result: the pooled cross-area classifier described here produces the shortlist performance of Section 2.1, the discrimination results of Sections 2.2 and 2.3, the worked examples of Section 2.5, and the conditional ranking of Section 2.6, with counts + cross-area precedent as its production feature set (Section 2.2).

### 3.4 Evaluation protocol

The primary discrimination analysis uses leave-one-focal-out (LOFO) evaluation across the 100-area panel. To score a held-out focal area in year Y, the model is trained only on the other areas using data strictly before Y, then applied to the held-out area at Y; features are standardised within each (area, cutoff) cell before fitting, and the abundant negatives are subsampled during training because the pooled cross-area base rate is well below one percent. During training, every positive was retained together with a uniform sample of twenty negatives per positive under a fixed seed. All held-out rows were scored and all reported metrics were calculated on the complete evaluation set, so the evaluation prevalence was not altered by training subsampling. For a logistic link, uniform down-sampling of one class multiplies the odds by a constant and so shifts only the intercept; for a gradient-boosted tree ensemble the training sample can in principle affect the learned ranking as well, so the ordering is not assumed to be preserved. The subsampling ratio was fixed in advance and is held constant across every configuration compared in this paper. Model and baselines are evaluated through a single pipeline on identical held-out rows.

Primary performance was evaluated using a temporally controlled historical replay designed to preserve the information available at each forecast origin. The protocol requires complete four-year outcome maturation, restricts evaluation to combinations still at risk of first clinical entry, and constructs partner precedent with the scored focal area held out. Concretely, its three controls are as follows. *Complete outcome-label maturation* applies on both sides of the split: on the evaluation side, a held-out row is scored only if its full four-year outcome window is observed within the focal area’s clinical record (Y+4 no later than that area’s last clinical-data year), so a recent cutoff cannot silently label an as-yet-unobserved future entry as a negative; and on the training side, a four-year gap (training cutoffs no later than Y−4) keeps the model from training on cutoffs whose own labels were themselves immature at training time. Enforcing maturity per focal area against its actual observed clinical extent is stricter than a single global data-freeze cut-off, because areas whose clinical record ends earlier are censored earlier. The *first-entry risk set* keeps only rows whose combination is not already in a trial as of Y, and excludes the in-trial indicator itself, so the target is genuinely a first entry rather than the re-identification of an existing one. The *leave-one-area-out precedent* construction ensures the scored area never contributes to its own cross-area precedent features (Section 3.2). A *relaxed-control sensitivity* (training on every prior cutoff earlier than Y, and scoring all held-out rows with the maturation and at-risk restrictions removed) quantifies what the controls cost (Section 2.3, Table 4). The temporally controlled replay is the protocol behind every primary table, the feature-group decomposition in Table 3, Appendix A and Figure 3.

This historical replay design is central. A score computed for year Y is a function only of literature, patent and trial records timestamped at or before year Y, under the source date fields of Section 3.1, so the study uses point-in-time construction rather than a conventional random split. The prediction target is first clinical entry within a 4-year horizon (entry in years Y+1 to Y+4, strictly after the cutoff). The panel is built from 2006; scoring cutoffs span 2010 to 2023, with the 2006 to 2009 build years supplying feature history but never scored, a floor applied at load before base rates or precedent are computed. Positivity is rare throughout: per-area positive rates on the evaluated rows range from under 0.1% to about 4%, with a median of 0.365%, and the pooled base rate under the temporally controlled replay is 0.50% (24.12 million scored combination-year rows across the 100 areas). This severe and variable class imbalance is why Section 2.3 reports precision-recall AUC alongside ROC-AUC.

### 3.5 Ground truth and baselines

For cross-area discrimination, each row is positive if the co-occurring keyword combination first enters a clinical trial within the forward 4-year window and negative otherwise. The negative class is therefore every burst-qualified co-occurring combination that did not enter a trial in that window, not a curated set of hard negatives. The main count-based baselines are cumulative co-occurring paper count (bibliometric count baseline), cumulative co-occurring patent count, and recent 3-year paper activity (recent-activity baseline), with the best of the three reported per area.

#### Enrichment

Precision-recall enrichment was calculated separately within each focal area as that area’s average precision divided by that area’s own event prevalence, after which the median ratio was reported across focal areas. This median-of-ratios is not interchangeable with the ratio of the median PR-AUC to the median prevalence, which is larger; only the per-area calculation is reported here.

#### First-alert precision (Sections 2.1, 2.2, 2.5 and 2.6)

The unified partner-level protocol scores an alert policy rather than the raw ranking, and every configuration compared in this paper (the core counts-and-precedent model, the full feature set, its ablations, and the zero-precedent conditional view) is evaluated under it identically. Keyword-row metrics, which count a partner once per synonym variant and once per cutoff, are not comparable with this deduplicated partner unit and are not used.

#### Metric

Keyword-level held-out scores are collapsed to a single partner-level value per (focal area, cutoff) cell by taking the maximum across resolved variants. Resolved within-vocabulary variants are collapsed to a single partner; aliases not linked by the cross-vocabulary resolution layer may remain separate partner identities. Each partner can occupy only one shortlist position; the top *k* unique partners of the cell are selected; a partner’s alert fires once, at its first selection; opening-year alerts are excluded. A hit is a first trial with the focal within the four years after the alert, the paper’s primary endpoint, and an alert enters the calculation only if that four-year window is fully observable, the same maturity logic as Section 3.4. When two score sets are compared, they are compared on identical cells: matched cutoffs, with a cell entering a depth only when both arms can fill it, so a missing score is never treated as a low score. Areas enter the analysis when at least 80% of the pooled panel’s partner vocabulary is resolvable against the outcome-timing tables and at least three matched cells exist; 97 of the 100 areas qualify.

### Null and multiplicity

The null is a matched random ranker: per cutoff, a uniform sample of k unique partners from the same cell with the same deduplication and censoring, drawn 2,000 times per area, the observed value’s percentile giving a one-sided p. Per-area p-values are reported nominally and under Benjamini-Hochberg at FDR 0.05. The panel-level omnibus compares the observed panel median against the null distribution of the panel median, one value per aligned draw. Enrichment against background divides each area’s precision by its own background four-year first-alert rate before taking the median across areas, with a 2,000-resample bootstrap CI on the median ratio. The zero-precedent conditional analysis of Section 2.6 applies the identical machinery with the candidate pool of every cell restricted to partners whose leave-one-area-out clinical-entry count is zero at that cutoff, and its null drawn from that restricted pool.

### Operating point

The twenty-partner review depth was an existing practical review depth and is used as the primary operating point. The other depths characterise the precision-coverage curve rather than serving as additional operating points.

### 3.6 Robustness checks

All robustness checks run on the temporally controlled replay of Section 3.4, on the full panel, through one pipeline on identical held-out rows; the relaxed-control sensitivity of Table 4 quantifies what the temporal controls cost. Four checks are applied. First, a fixed-score label permutation confirms that the reported metric and the focal-area aggregation are centred at chance under within-area label exchange (Section 2.3). Second, the feature ablation of Table 3 runs every feature group through the identical protocol, isolating the precedent contribution from the burst-and-count contribution. Third, the event study of Section 2.4 verifies temporally that the clinical-source channel carries no signal before a pair’s first trial exists, so the pre-trial signal is textual. Fourth, the unified first-alert analysis of Section 2.1 acts as the operational check: it evaluates the ranking as an alert policy on deduplicated partners, under a matched random null, which is the harder and operationally relevant question. Given the severe class imbalance, we also report precision-recall AUC alongside ROC-AUC. These checks do not prove the absence of every possible artefact, but they test the most plausible leakage and interpretation concerns.

### 3.7 Validation controls and reproducibility

Historical replay requires that every model input, training outcome and candidate-eligibility decision reflect information available at the scoring cutoff. The evaluation therefore applies the three temporal controls of Section 3.4 throughout: complete four-year outcome maturation on both training and evaluation rows, a first-entry risk set excluding combinations already in trials at the cutoff, and leave-one-area-out construction of partner precedent. The headline is robust to control strictness (imposing the full controls against the relaxed-control sensitivity moves the median AUC by 0.011, Table 4), and the panel’s high cross-area partner recurrence (Section 2.3) is a plausible reason the sensitivity is modest, since information about a partner persists across many areas under any one restriction.

#### Reproducibility

Analysis code, model configuration and input-data provenance were version controlled, and structural checks verified the expected row counts, outcomes and held-out assignments.

## 4. Related work

Table 8 positions this work among the closest adjacent systems. Among them we did not identify one that combines this work’s unit (a specific focal-partner combination), endpoint (first clinical entry within four years) and temporal design (a historical replay with the scored area excluded from training); the subsections that follow give the fuller account of each literature.

**Table 8.**
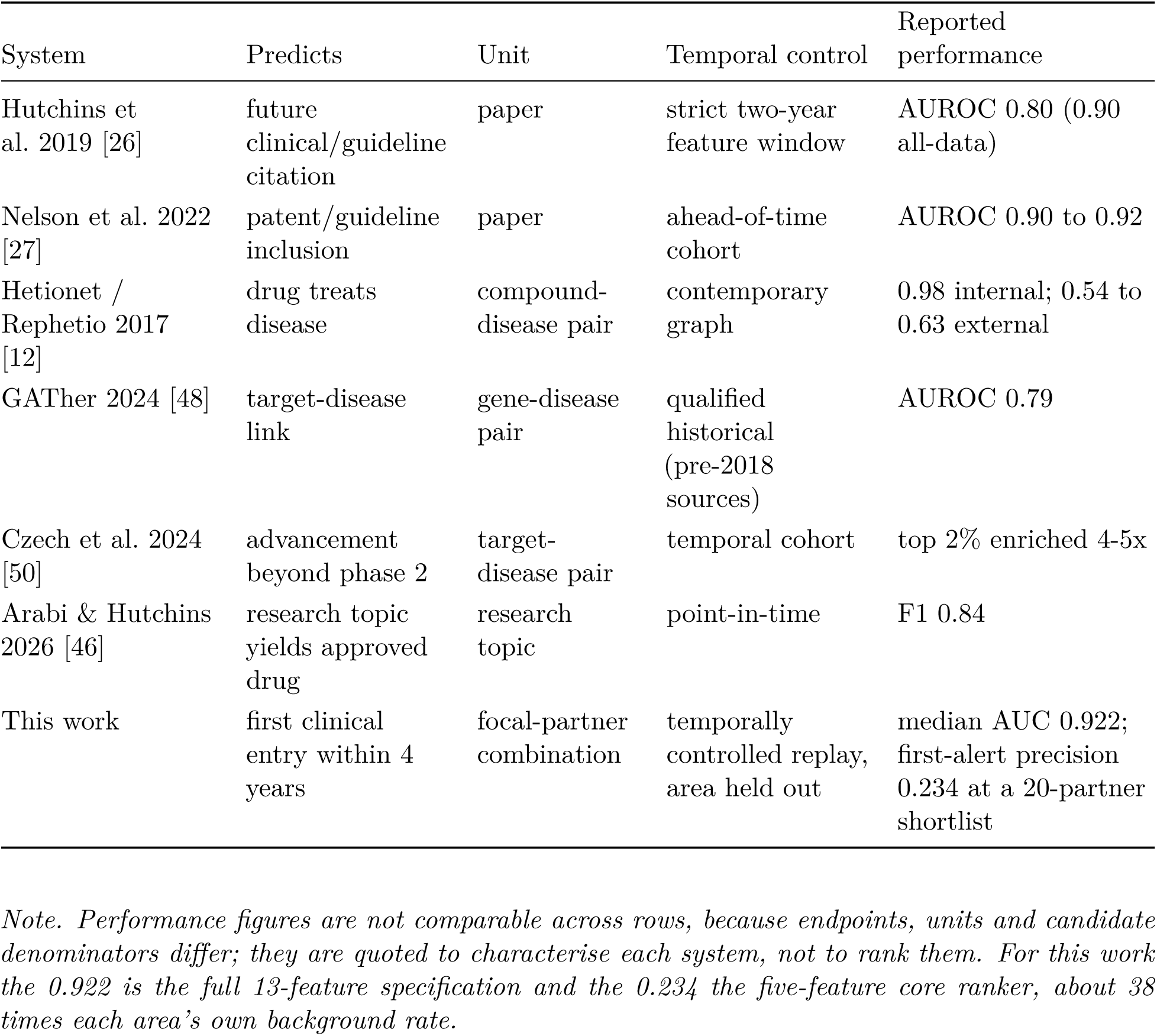
The closest adjacent systems, by prediction target, unit and temporal control.

### 4.1 Literature-based discovery and hypothesis generation

Literature-based discovery began with Swanson’s demonstrations that disjoint literatures can imply clinically relevant hypotheses without any single paper stating the connection explicitly [4]. The field later broadened from simple ABC models to ranking methods, semantic predication systems, and graph-based recovery of latent links [7–9]. Reviews and scientometric mappings show a progression from co-occurrence-driven methods toward semantic, embedding, and explainability-oriented systems [5, 6, 10, 28]. A parallel line represents the literature not by explicit co-occurrence but by distributed embeddings: unsupervised word embeddings have been shown to anticipate later scientific discoveries [17], and biomedical-specific representations (subword- and MeSH-aware word embeddings, transformer language models, and concept embeddings) encode latent structure in the same corpora we draw on [18–20]. These representations are complementary to the explicit, point-in-time burst features studied here, which we favour for their auditability under strict historical replay. More recently, foundation models and large language models have been advanced as general engines for scientific discovery [45]. We take a complementary approach for this triage layer, using explicit, interpretable features so that each prediction can be traced to a specific acceleration in the historical evidence record.

### 4.2 Knowledge graphs and link prediction for drug repurposing

Knowledge graphs provide a richer representation of biomedical entities and relations than term co-occurrence alone. Hetionet-style integration of curated and observational sources has been used to prioritise repurposing opportunities [12], and later work extended this direction with graph embeddings, graph neural networks, and large integrated target-prioritisation platforms [14–16, 29–31]. These approaches can represent heterogeneous evidence and complex multi-hop structure, but temporal validation remains diffiicult when source graphs are assembled from knowledge accumulated over many years. The consequence is measurable: Project Rephetio’s internal evaluation reports AUROC near 0.98, while external evaluation of the same system against independent indication sets falls to 0.54 and 0.63 [12]: a graph can separate known positives from sampled negatives under its own construction while failing to transport beyond the training frame. The closest genuinely historical comparator at the target level is GATher, a graph-attention model trained on sources dated before 2018 and evaluated on clinical development through 2024, reporting AUROC 0.79 for positive-effiicacy outcomes and 14.1% precision among its top 200 target-disease predictions that later enter trials [48]. That is a target-disease unit and a downstream effiicacy endpoint rather than first entry of a specific focal-partner combination, but it supports the pattern this paper’s own controls show: strict or qualified historical designs report lower and more credible performance than present-day graph holdouts.

### 4.3 Bibliometric prediction of translation and emerging topics

Burst detection and research-front analysis have long been used to identify emerging topics [21–23]. Later studies asked not just whether a term is growing, but whether its future trajectory can be forecast [24]. In translational science, bibliometric and text-mining approaches have been used to predict whether publications will be cited by future trials or guidelines, to quantify translational movement across research stages, and to trace links among funding, publications, patents, and approved drugs [26, 27, 32, 33]. The present work differs in that its unit is not an individual paper but a target-pair or keyword-combination signal, and its endpoint is clinical entry within a fixed forward horizon. A distinct but sequential line predicts clinical success itself (whether a trial or drug will ultimately win approval) from trial-level features [44]; that task begins where ours ends, at trial entry, so the two are complementary stages of one pipeline rather than competing approaches to the same question. Most directly related, a recent preprint forecasts which research topics will yield FDA-approved drugs from citation-graph features alone (citation patterns, publication content, and researcher migration into emerging topics), reporting an F1 of 0.84 and flagging a majority of eventual drug targets years before phase 2 [46]. Like the present study, that method is evaluated point-in-time. Its predictors are derived from citation-graph features without using trial records, whereas Intangia’s pooled model incorporates point-in-time clinical precedent from other focal contexts; and it targets eventual approval at research-topic level, a further-downstream and coarser endpoint than the first clinical entry of a specific focal-partner combination.

### 4.4 Cross-area clinical precedent and target maturity

The central empirical finding of this paper, that cross-area clinical precedent dominates prediction of first entry, sits inside an established literature on how drug development clusters around validated targets. Swinney and Anthony’s analysis of 1999 to 2008 approvals found only 75 of 259 new molecular entities were first-in-class, leaving most approvals in precedented or follower categories [49]; prior trials on a target reduce uncertainty not only about biology but about tractability, safety liabilities, assay infrastructure and investor credibility, so precedent is a portfolio signal as much as a biological one. The closest quantitative comparator is Czech et al.’s clinical advancement forecasting: among 9,010 target-disease pairs entering phase 2 between 2016 and 2022, 94% involved targets with prior phase-2-or-higher trials, and a multievidence model identified a top 2% of pairs four- to fivefold more likely to advance beyond phase 2 [50]. McNamee et al. report that targets entering trials after an established point of technology maturation develop substantially faster than less mature ones [51]. The settings are not directly comparable (Czech et al.’s top 2% of target-disease pairs is four-to fivefold more likely to advance beyond phase 2 [50], while our twenty-partner shortlist is about 38-fold enriched over background at the earlier and rarer endpoint of first clinical entry, Section 2.1), but both show the same qualitative fact: evidence-and-precedent models concentrate future clinical activity strongly. Against this literature the present contribution is not the observation that precedent matters, which is well documented, but its leave-one-area-out quantification for first clinical entry of focal-partner combinations, under a replay in which the scored area never contributes to its own precedent. The same literature frames the model’s measured boundary: a precedent-dominated ranker performs best where development follows known target classes. The zero-precedent stratum of Section 2.6, related to though not the same as the first-in-class setting in which the field’s opportunity arguments for underexplored but tractable targets are made [52], is served by the conditional emerging-opportunities view at a lower yield.

### 4.5 Drug combination prediction in oncology

Computational prediction of drug combinations has largely emphasised synergy, response, or toxicity rather than historical anticipation of clinical entry. Representative approaches include deep-learning models for anti-cancer drug synergy, interpretable response predictors, and curated combination data resources [34–37]. Those methods address a different question from the one studied here: whether a candidate combination is likely to be effective in a biological or screening context, rather than whether the preclinical record already contains a signal of translation.

### 4.6 Patent analysis for competitive intelligence

Patent corpora have been mined to characterise technological trends, identify emerging biomedical concepts, and assess commercial interest across diseases and target classes [38–40]. In immuno-oncology specifically, combined journal-and-patent analyses have been used to map emerging targets and therapeutic concepts at scale [40]. The worked examples of Section 2.5 bear this out: in both areas, a majority of the replayed true-positive alerts carried more co-occurring patent than paper evidence at the alert cutoff, most visibly CD47’s CTLA-4 alert, which rested on 107 co-occurring patents against 5 papers four years before the first registered trial pairing the two.

## 5. Discussion

What the results change is the economics of looking. Portfolio and business-development teams do not suffer from a shortage of candidate pairings; they suffer from a review budget that is fixed while the landscape is not. A twenty-name shortlist in which roughly a quarter of the names go on to enter the clinic converts an open-ended scanning problem into a bounded weekly review, and it does so early enough that the finding can still change a decision rather than merely explain one. That the same ranking holds across a hundred areas spanning oncology to rare disease means the capability is a property of the evidence record rather than of any one therapeutic story, so a team can point it at a new area without first asking whether that area is the kind where it works.

Why it works is legible in the feature decomposition, and the division of labour is the architecture. Cumulative evidence counts and cross-area clinical precedent carry the ranking: partners already pursued clinically in one context are more likely to be pursued in another, which is what the target-maturity literature predicts (94% of recent phase-2 entries involve targets already in phase-2-or-higher trials [50]), and the contribution here is its leave-one-area-out quantification for focal-partner entry (Section 4.4). Burst detection carries momentum and explanation: it identifies where activity is accelerating, which sources are driving the acceleration, and when it began, and it attaches that time-stamped rationale to every recommendation. Its measured contribution to the ranking itself is not material (Section 2.4), and the product does not ask it to be: the burst channel is what distinguishes an emerging opportunity from a static incumbent, often moving before the ranking promotes a candidate, and it is what a reviewer interrogates when deciding why now.

Relative to knowledge graph and graph neural network methods, the approaches are complementary: KG and GNN systems represent mechanistic richness and multi-hop biology more naturally [12, 14, 15], but historically faithful temporal validation is hard once a graph aggregates years of curated knowledge. The advantage here is auditability under explicit temporal controls and a temporally controlled point-in-time replay. Each signal traces to a source-specific acceleration in the pre-cutoff record, and the burst signature is itself a structured hypothesis (where the acceleration originated, when it began, and whether it is context-specific), which is easier to hand to downstream mechanistic or expert review than a scalar probability.

The subsections below set out what this capability changes for the teams that use it. Section 6 sets out where the evidence stops.

### 5.1 Faster landscape triage

A team choosing where to commit early experimental or analyst effort faces a candidate space far larger than it can evaluate: typically thousands of scored candidates per focal area in this panel. The useful question is not whether a ranking is perfect but whether it concentrates future entrants near the top, and this one does: at the twenty-partner review depth, roughly one in four shortlisted partners entered the focal clinical context within four years, about 38 times the area’s own background rate (Section 2.1). A landscape too large to survey by hand is reduced to a reviewable top twenty, with the evidence behind each entry attached to it, and the depth schedule of Table 1 lets a team trade precision for coverage explicitly.

### 5.2 Better allocation of diligence

Early triage is a high-volume, high-cost screening problem: every candidate advanced to mechanistic or experimental evaluation consumes analyst time or laboratory resource. Because a twenty-partner shortlist converts at roughly one in four rather than the background’s well under one in a hundred, recovering a given number of genuine entrants takes far fewer downstream evaluations than an unranked or chance-level process would require, and the precision-versus-depth curve of Figure 2 prices the alternative depths. The shortlist arrives as an auditable, time-stamped hypothesis rather than a single risk score, so the reviewer can interrogate it.

### 5.3 Earlier strategic positioning

Recommendations are generated before clinical entry, so there is a real decision window in which to do the work that has to happen early: testing the biology, developing differentiated evidence, assessing intellectual-property space, evaluating build-versus-partner options, and positioning a programme before its window closes.

The scope of this is broader than the combination-therapy framing might suggest. The method operates on any focal-partner pairing (Box 1), and the cross-area panel spans both drug targets and disease indications, so the framework also applies to mechanism-by-indication opportunities, the common monotherapy case in which the larger share of drug development sits. In less-precedented settings, the Emerging Opportunities view of Section 2.6 provides a lower-confidence lens before substantial clinical precedent has accumulated. The PD-1 combination case is the worked example; the breadth result (Section 2.2) is the evidence that the capability generalises across contexts, and first-in-class prediction as such is not validated here.

### 5.4 Consistency and institutional memory

Competitive-intelligence and research teams already track emerging programmes through conference activity, expert networks, patent monitoring and manual literature review. Intangia systematises those cues: it applies the same evidence criteria across every focal area, identifies changes on a consistent basis rather than as attention allows, and preserves the historical provenance of each flag so the rationale can be replayed at the date it was made. That last property is what turns triage into institutional memory. A judgement made three years ago can be re-examined on the evidence that actually supported it, rather than reconstructed from recollection or lost when the analyst who made it moves on.

### 5.5 Evaluation transparency

Any system claiming to predict clinical translation from the preclinical record faces the same temporal-validity requirements this study imposes on itself: point-in-time feature construction, held-out-area evaluation, matured outcome labels, and performance reported as a deduplicated alert policy against a matched random benchmark rather than only as a ranking metric. Adjacent fields have moved the same way (medicinal chemistry’s SIMPD algorithm generates simulated time splits precisely because random splits flatter models that will face temporal distribution shift in use [53]), and reporting standards such as TRIPOD+AI point in the same direction [47]. Because the method under evaluation is also a commercial product, we report the permutation control, feature ablation, control-strictness sensitivity and matched-random operational benchmark in full, and an anonymised validation package is available for independent technical review (Data and code availability).

## 6. Limitations

The claims above are bounded in three stateable ways, with the technical measurement notes gathered at the end. Each is set out once here; the sections that depend on one refer back rather than restating it.

### The endpoint is where development begins, not whether it succeeds

The model identifies where clinical activity is likely to occur next; it does not determine whether the resulting programme will work, and historical success rates from first-in-human testing to approval are low and strongly phase-dependent [43]. Biological causality, technical feasibility and commercial value are assessed downstream of the layer evaluated here. Combination-level discrimination is measured against every burst-qualified co-occurrence that did not enter trials, not a curated set, so the scope boundary is the endpoint, not the denominator.

#### Timing is reported descriptively; demonstrating timing skill is the next validation milestone

Lead time throughout this paper is the observed interval from first alert to first trial. A ranking that promotes a partner early and keeps it promoted accumulates lead time by persistence, so the observed intervals establish that alerts land materially before entry, not that the system times entries better than a comparably persistent alternative would. What is established quantitatively is selection: at every review depth, shortlisted partners enter at many times the matched-random and background rates (Section 2.1). Every result here is retrospective; whether organisations convert the signal into demonstrably earlier, better portfolio decisions is the prospective question Section 7 sets out.

#### The model is strongest where clinical precedent exists

The median area shares 99.8% of its partners with at least one other area (minimum 97.5%), and the headline results are therefore predictions across focal areas for such partners. Among partners with no cross-area precedent, the conditional view of Section 2.6 is enriched relative to matched random ranking but at substantially lower absolute yield, and zero precedent within this panel is not synonymous with first-in-class status.

#### Measurement notes

Three construction details bound specific numbers rather than the claims. (i) The two worked-example areas were selected post hoc as familiar illustrative examples from the panel and are not used to estimate performance in an unseen focal area; the performance estimates are the panel-wide analyses of Sections 2.1 and 2.2. (ii) Sponsor and assignee fields are not yet in the feature set, so field-wide convergence is not separated from a single organisation documenting a programme it already intends to advance. (iii) Patents are counted per publication rather than per family, which inflates patent volume and smears its timing (Section 3.1).

## 7. Future validation

### Prospective replay

Freeze current rankings across a pre-specified set of focal areas and report clinical-entry outcomes at fixed future checkpoints. The same exercise is the right place to formalise the which-versus-when distinction this paper draws informally: survival methodology (horizon-specific calibration, time-dependent discrimination and prediction-accuracy indices for censored outcomes [54, 55]) would let timing skill be claimed or refuted directly rather than reported descriptively.

### Low-precedent extension

The zero-precedent setting is measured directly: the conditional view of Section 2.6 ranks eventual entrants above the matched-random benchmark among their zero-precedent peers (38% of the 170 unique entrants reach the conditional top twenty before entry) at an absolute precision roughly an order of magnitude below the core ranking. That boundary is a consequence of the model’s deliberate strength: it transfers clinical precedent across contexts, and here there is little to transfer. Raising the yield of the emerging-opportunities view, in the setting closest to first-in-class discovery where an early signal would be worth most [52], calls for its own baseline and feature family, and is the highest-value extension on this roadmap. The exploratory acceleration regimes of Section 2.4 are the first candidates to test as that record accumulates.

### First-posted-date sensitivity

Re-slice the clinical series on the registry first-posted date rather than the first-submission timestamp, and re-run the replay. This would establish that every clinical record used at a scoring date was publicly observable at that date, closing the gap noted in Section 3.1. It is the one temporal control this study could not run from its own inputs: the analysis consumes a pre-aggregated co-occurrence extract in which the clinical leg is already year-keyed on first submission, so the sensitivity requires rebuilding that leg from the raw registry upstream, not a reanalysis of the extract. It matters disproportionately because cross-area clinical precedent is the load-bearing predictor (Section 2.2).

### Sponsor-controlled analysis

Separate broader field convergence from within-company publication, patent and trial activity, using the sponsor and assignee fields availh3able in the underlying source records.

### Pre-registered panel rules

The fifty-positive reporting floor of Section 3.1 is applied at analysis time; fixing panel-inclusion thresholds *before* a panel is drawn, as part of a registered prospective design, is the cleaner standard the prospective replay above should adopt.

### A non-trivial external baseline

Every baseline in this paper is count-based. The natural stronger comparator is a target-maturity model in the style of Czech et al.’s clinical advancement forecasting [50] or an Open Targets-style composite [16], reconstructed at each forecast origin from archived evidence and mapped to focal-partner entry. Beating naive counting is established here; beating an archived maturity score is the next bar, and Section 4.4’s comparison suggests it is the right one, since that is the strongest signal family this model itself relies on.

### External and sparse-area validation

Extend the panel to emerging areas with limited evidence and support independent reproduction using an anonymised validation package.

## 8. Conclusion

This study validates Intangia’s point-in-time triage layer as a working shortlist engine. Across 100 focal areas and 24.12 million historically scored combination-years, a single pooled cross-area recommender, ranking on cumulative evidence counts and leave-one-area-out clinical precedent, delivered four-year first-alert precision of 0.234 at a twenty-partner review depth against a matched-random median of 0.008: roughly one in four shortlisted partners subsequently entered the focal clinical context within four years, about 38 times each area’s own background rate, with 94 of 97 areas surviving false-discovery control and a panel-level permutation at empirical *p* ≤ 0.0005. The ranking generalises (the full 13-feature specification reaches a median leave-one-focal-out AUC of 0.922 and the five-feature core ranker 0.915, with no area below chance and every area beating its own best count-based baseline, each area scored by a model that never saw it), and shortlisted entrants are anticipated with a median observed lead of two years.

Around that engine, burst detection supplies what a scalar ranking cannot: time-stamped, source-specific evidence of momentum, attached to every recommendation and frequently visible before the ranking promotes a candidate. A conditional view of the same landscape ranks zero-precedent candidates against one another, enriched relative to matched random ranking, at a deliberately lower-yield setting that supports an emerging-opportunities lens rather than a second model. The worked examples in PD-1 combination immunotherapy and CTLA-4 show the product behaviour concretely: of the replayed shortlists’ 82 and 53 alerts, 59 and 19 partners entered a trial with the focal within four years, each with the dated record that produced the alert.

The endpoint is first clinical entry rather than therapeutic success or portfolio value, and lead time is reported descriptively. Within that defined scope, the evidence supports Intangia’s triage layer as a credible early-warning and prioritisation capability for drug-development decision-making. Prospective validation is the next stage.

## About Intangia

The triage layer evaluated here is one component of the Intangia platform, which supports therapeutic-area landscaping, target prioritisation, portfolio strategy and business-development assessment. This preprint reports the retrospective validation of the triage layer specifically; the downstream causal-evidence and scoring layers shown in Figure 1 are evaluated separately. Enquiries about the platform, about the validation package described below, or about evaluating the method on a specific therapeutic area, are welcome.

## Data and code availability

The underlying co-occurrence tables contain proprietary Intangia transformations and licensed source data and cannot be released publicly. For technical review, Intangia can provide an anonymised validation package containing model features, outcome labels, focal-area and cutoff identifiers, the model configuration, and suffiicient documentation to reproduce the reported discrimination metrics. Requests should be directed to the corresponding author.

## Acknowledgements

We thank Pascal Deschatelets (independent biotech advisor) for helpful comments on a draft.

## Appendix A. Per-area discrimination, all evaluable focal areas

The summary statistics in Table 2 (median AUC 0.922, worst-case 0.818, no inverted areas) and the ranked distribution in Figure 3 are reported here in full, one row per evaluable focal area, so the breadth claim can be checked area by area rather than in aggregate. Each row gives the full 13-feature specification’s temporally controlled leave-one-focal-out clinical-entry AUC at the 4-year horizon, that area’s best count-based bibliometric baseline on the same held-out rows, and the lift of the model over that baseline; areas are ordered from lowest to highest model AUC. All 100 areas are named. The number of scored rows and positives per area is included so that areas with thin positive counts, where the AUC is noisier, are visible. Two areas fall below the fifty-positive floor adopted in Section 3.1: CACNA1H with 11 positives and an AUC of 0.941, and RBX1 with 39 and 0.961. Both sit above the panel median, so they bias it upward rather than down; removing them moves the median from 0.922 to 0.920 and leaves the minimum unchanged.

**Table A1.** Per-area leave-one-focal-out clinical-entry AUC under the temporally controlled replay (4-year horizon) for all 100 evaluable focal areas, ranked. Median model AUC 0.922; minimum 0.818; areas below 0.50: 0.

| # | Focal area | Rows<br>(n) | Positives | Model<br>AUC | Best<br>baseline | Lift |
| --- | --- | --- | --- | --- | --- | --- |
| 1 | Non-small cell lung carcinoma | 354,732 | 3424 | 0.818 | 0.731 | +0.087 |
| 2 | Somatostatin receptor 5 | 30,398 | 152 | 0.819 | 0.689 | +0.130 |
| 3 | Head and neck squamous cell carcinoma | 159,784 | 1850 | 0.828 | 0.724 | +0.104 |
| 4 | Triple-negative breast carcinoma | 553,060 | 4096 | 0.829 | 0.707 | +0.122 |
| 5 | Phospholipase A2 group IIA | 437,721 | 3291 | 0.836 | 0.744 | +0.092 |
| 6 | PD-1 | 331,647 | 4040 | 0.840 | 0.713 | +0.127 |
| 7 | Ovarian cancer | 426,083 | 3035 | 0.853 | 0.749 | +0.104 |
| 8 | Malignant pancreatic neoplasm | 388,672 | 3173 | 0.862 | 0.757 | +0.105 |
| 9 | Steroid 5 alpha-reductase 2 | 83,055 | 126 | 0.869 | 0.766 | +0.103 |
| 10 | Gastroesophageal junction<br>adenocarcinoma | 9,971 | 351 | 0.875 | 0.694 | +0.180 |
| 11 | Non-Hodgkin lymphoma | 281,828 | 1329 | 0.875 | 0.760 | +0.115 |
| 12 | Melanoma | 562,018 | 2788 | 0.876 | 0.769 | +0.107 |
| 13 | Endometrial cancer | 263,569 | 2043 | 0.882 | 0.776 | +0.106 |
| 14 | Interleukin 12B | 77,852 | 180 | 0.886 | 0.686 | +0.201 |
| 15 | Fms related receptor tyrosine kinase 3 | 246,485 | 736 | 0.887 | 0.712 | +0.175 |
| 16 | Fallopian tube cancer | 59,155 | 579 | 0.888 | 0.758 | +0.130 |
| 17 | Gastric cancer | 282,071 | 2096 | 0.889 | 0.752 | +0.137 |
| 18 | Prostaglandin-endoperoxide synthase 2 | 368,826 | 1985 | 0.889 | 0.735 | +0.155 |
| 19 | Sodium voltage-gated channel alpha<br>subunit 1 | 83,744 | 318 | 0.890 | 0.695 | +0.195 |
| 20 | KIT proto-oncogene, receptor tyrosine<br>kinase | 226,962 | 1090 | 0.890 | 0.706 | +0.184 |
| 21 | Esophageal cancer | 246,940 | 1384 | 0.891 | 0.776 | +0.115 |
| 22 | Multiple myeloma | 437,644 | 1401 | 0.893 | 0.756 | +0.136 |
| 23 | Fms related receptor tyrosine kinase 1 | 285,994 | 1057 | 0.894 | 0.760 | +0.134 |
| 24 | Kinase insert domain receptor | 389,282 | 1415 | 0.897 | 0.748 | +0.149 |
| 25 | Follicular lymphoma | 139,864 | 863 | 0.897 | 0.767 | +0.130 |
| 26 | Fms related receptor tyrosine kinase 4 | 239,231 | 821 | 0.898 | 0.732 | +0.166 |
| 27 | Androgen receptor | 204,465 | 1285 | 0.901 | 0.714 | +0.187 |
| 28 | 5-hydroxytryptamine receptor 2A | 158,082 | 656 | 0.902 | 0.764 | +0.137 |
| 29 | Acute lymphoblastic leukaemia | 316,922 | 814 | 0.902 | 0.785 | +0.117 |
| 30 | Diabetic neuropathy | 145,833 | 222 | 0.902 | 0.693 | +0.209 |
| 31 | Transient receptor potential cation<br>channel subfamily V member 1 | 104,166 | 523 | 0.903 | 0.680 | +0.223 |
| 32 | TNF-alpha | 586,582 | 1061 | 0.903 | 0.767 | +0.136 |
| 33 | Sodium voltage-gated channel alpha<br>subunit 5 | 150,027 | 532 | 0.904 | 0.716 | +0.189 |
| 34 | Urothelial carcinoma | 73,590 | 1200 | 0.905 | 0.716 | +0.189 |
| 35 | Histamine receptor H2 | 37,606 | 308 | 0.906 | 0.675 | +0.231 |
| 36 | Psoriatic arthritis | 171,308 | 397 | 0.906 | 0.816 | +0.090 |
| 37 | Glioblastoma multiforme | 215,540 | 1121 | 0.907 | 0.736 | +0.171 |
| 38 | Opioid receptor mu 1 | 172,646 | 574 | 0.909 | 0.724 | +0.185 |
| 39 | Cholinergic receptor muscarinic 3 | 71,464 | 256 | 0.909 | 0.717 | +0.192 |
| 40 | GLP-1R | 174,666 | 753 | 0.909 | 0.771 | +0.138 |
| 41 | PD-L1 (CD274) | 306,936 | 2414 | 0.911 | 0.746 | +0.165 |
| 42 | EGFR | 546,667 | 1377 | 0.912 | 0.733 | +0.179 |
| 43 | PPAR-gamma | 240,508 | 424 | 0.913 | 0.693 | +0.220 |
| 44 | VEGF-A | 477,990 | 1584 | 0.914 | 0.741 | +0.173 |
| 45 | Psoriasis | 351,879 | 716 | 0.914 | 0.837 | +0.077 |
| 46 | Prostaglandin-endoperoxide synthase 1 | 474,602 | 425 | 0.914 | 0.796 | +0.118 |
| 47 | Mesothelioma | 214,348 | 1162 | 0.915 | 0.829 | +0.086 |
| 48 | Raf-1 proto-oncogene, serine/threonine kinase | 181,818 | 741 | 0.918 | 0.712 | +0.206 |
| 49 | Solute carrier family 6 member 2 | 162,419 | 470 | 0.919 | 0.786 | +0.133 |
| 50 | HER2 (ERBB2) | 406,686 | 1126 | 0.921 | 0.732 | +0.189 |
| 51 | Cereblon | 25,922 | 84 | 0.922 | 0.709 | +0.213 |
| 52 | CD19 | 318,036 | 929 | 0.923 | 0.811 | +0.113 |
| 53 | KRAS G12C | 3,444 | 137 | 0.924 | 0.773 | +0.151 |
| 54 | Janus kinase 1 | 190,135 | 1317 | 0.925 | 0.698 | +0.227 |
| 55 | Biliary tract cancer | 71,296 | 632 | 0.926 | 0.685 | +0.241 |
| 56 | Calcium voltage-gated channel subunit alpha1 C | 23,218 | 72 | 0.927 | 0.826 | +0.101 |
| 57 | Toll like receptor 7 | 172,630 | 1002 | 0.927 | 0.719 | +0.208 |
| 58 | Sarcopenia | 75,772 | 133 | 0.927 | 0.811 | +0.116 |
| 59 | CTLA-4 | 79,019 | 671 | 0.928 | 0.639 | +0.288 |
| 60 | Opioid receptor kappa 1 | 114,186 | 283 | 0.929 | 0.741 | +0.188 |
| 61 | Tyrosine kinase 2 | 147,481 | 788 | 0.929 | 0.721 | +0.208 |
| 62 | Dopamine receptor D2 | 112,821 | 493 | 0.929 | 0.722 | +0.207 |
| 63 | Coagulation factor X | 250,766 | 524 | 0.930 | 0.827 | +0.103 |
| 64 | IL-17A | 258,142 | 672 | 0.934 | 0.805 | +0.130 |
| 65 | 5-hydroxytryptamine receptor 3A | 110,407 | 580 | 0.935 | 0.760 | +0.175 |
| 66 | Glutamate ionotropic receptor NMDA type subunit 2A | 119,775 | 164 | 0.935 | 0.774 | +0.161 |
| 67 | MEK1 (MAP2K1) | 229,644 | 870 | 0.938 | 0.723 | +0.215 |
| 68 | Janus kinase 3 | 187,105 | 1150 | 0.939 | 0.702 | +0.236 |
| 69 | DNMT1 | 199,374 | 861 | 0.939 | 0.679 | +0.260 |
| 70 | BCMA | 159,146 | 431 | 0.939 | 0.826 | +0.113 |
| 71 | Cytochrome P450 family 3 subfamily A member 7 | 67,881 | 361 | 0.939 | 0.665 | +0.274 |
| 72 | Phosphodiesterase 5A | 133,100 | 601 | 0.939 | 0.694 | +0.245 |
| 73 | Cholinergic receptor muscarinic 2 | 58,100 | 97 | 0.940 | 0.657 | +0.283 |
| 74 | Gamma-aminobutyric acid type A receptor subunit theta | 196,138 | 386 | 0.941 | 0.756 | +0.185 |
| 75 | Calcium voltage-gated channel subunit alpha1 H | 7,121 | 11 | 0.941 | 0.790 | +0.150 |
| 76 | CNS neoplasm | 121,979 | 461 | 0.941 | 0.768 | +0.173 |
| 77 | ABL proto-oncogene 1, non-receptor tyrosine kinase | 285,367 | 1025 | 0.942 | 0.718 | +0.223 |
| 78 | PARP1 | 287,724 | 1249 | 0.942 | 0.735 | +0.207 |
| 79 | DNA methyltransferase 3 alpha | 181,356 | 850 | 0.942 | 0.688 | +0.254 |
| 80 | 5-hydroxytryptamine receptor 2C | 108,944 | 425 | 0.943 | 0.735 | +0.208 |
| 81 | Platelet derived growth factor receptor alpha | 212,307 | 643 | 0.943 | 0.778 | +0.166 |
| 82 | Gonadotropin releasing hormone receptor | 87,658 | 225 | 0.946 | 0.710 | +0.236 |
| 83 | Solute carrier family 6 member 4 | 126,390 | 474 | 0.948 | 0.700 | +0.247 |
| 84 | Carbonic anhydrase 4 | 104,957 | 150 | 0.948 | 0.719 | +0.229 |
| 85 | Interferon alpha and beta receptor subunit 1 | 138,826 | 208 | 0.951 | 0.668 | +0.283 |
| 86 | Glutamate ionotropic receptor NMDA type subunit 1 | 236,133 | 314 | 0.951 | 0.738 | +0.213 |
| 87 | Fibroblast growth factor receptor 1 | 221,551 | 763 | 0.952 | 0.756 | +0.196 |
| 88 | CDK4 | 230,507 | 1185 | 0.953 | 0.763 | +0.189 |
| 89 | Glutamate ionotropic receptor NMDA type subunit 2B | 157,945 | 260 | 0.954 | 0.714 | +0.240 |
| 90 | Dopamine receptor D4 | 46,217 | 167 | 0.955 | 0.744 | +0.211 |
| 91 | CGRP receptor (CALCRL) | 92,360 | 153 | 0.955 | 0.774 | +0.181 |
| 92 | Fibroblast growth factor receptor 2 | 281,595 | 927 | 0.960 | 0.710 | +0.250 |
| 93 | Phosphodiesterase 6B | 41,906 | 79 | 0.960 | 0.611 | +0.349 |
| 94 | Ring-box 1 | 35,345 | 39 | 0.961 | 0.736 | +0.225 |
| 95 | Interleukin 2 receptor subunit beta | 206,443 | 574 | 0.962 | 0.765 | +0.197 |
| 96 | PCSK9 | 128,306 | 288 | 0.963 | 0.799 | +0.164 |
| 97 | Dopamine receptor D3 | 90,199 | 331 | 0.969 | 0.715 | +0.254 |
| 98 | CD3 (CD3E) | 162,772 | 290 | 0.970 | 0.696 | +0.274 |
| 99 | SIRP-alpha | 168,320 | 356 | 0.977 | 0.877 | +0.100 |
| 100 | Interleukin 2 receptor subunit gamma | 60,358 | 144 | 0.986 | 0.712 | +0.273 |

## Domain composition

So the breadth claim can be checked for therapeutic-area diversity rather than a hidden concentration in one field, Table A2 gives the coarse therapeutic domain of all 100 areas together with each domain’s median and minimum AUC. The panel is oncology-weighted at two areas in five, but is not oncology-only: the remaining sixty span six further domains. Oncology is also the weakest of the large strata, at a median of 0.900 against 0.935 for neuroscience and psychiatry and 0.933 for cardiometabolic areas, and it contains the panel minimum; only the four-area endocrine and genitourinary group sits lower, at 0.885. That ordering is the density finding of Section 2.2 restated by domain: oncology areas are the most heavily co-published, so co-occurrence carries least information about any specific pairing within them. The panel’s headline is therefore not being carried by its largest domain.

**Table A2.** Therapeutic-domain composition of the 100 focal areas.

| Domain | Areas | Median AUC | Minimum AUC |
| --- | --- | --- | --- |
| Oncology | 40 | 0.900 | 0.818 |
| Neuroscience / psychiatry | 16 | 0.935 | 0.890 |
| Immunology / inflammation | 16 | 0.926 | 0.836 |
| Haematology / rare disease / other | 14 | 0.927 | 0.906 |
| Cardiometabolic | 8 | 0.933 | 0.902 |
| Endocrine / genitourinary | 4 | 0.885 | 0.819 |
| Ophthalmology | 2 | 0.954 | 0.948 |
| Total | 100 | 0.922 | 0.818 |

